# FUN-LDA: A LATENT DIRICHLET ALLOCATION MODEL FOR PREDICTING TISSUE-SPECIFIC FUNCTIONAL EFFECTS OF NONCODING VARIATION

**DOI:** 10.1101/069229

**Authors:** Daniel Backenroth, Zihuai He, Krzysztof Kiryluk, Valentina Boeva, Lynn Pethukova, Ekta Khurana, Angela Christiano, Joseph D. Buxbaum, Iuliana Ionita-Laza

## Abstract

We describe here a new method based on a latent Dirichlet allocation model for predicting functional effects of noncoding genetic variants in a cell type and tissue specific way (FUN-LDA) by integrating diverse epigenetic annotations for specific cell types and tissues from large scale epigenomics projects such as ENCODE and Roadmap Epigenomics. Using this unsupervised approach we predict tissue-specific functional effects for every position in the human genome. We demonstrate the usefulness of our predictions using several validation experiments. Using eQTL data from several sources, including the Genotype-Tissue Expression project, the Geuvadis project and TwinsUK cohort, we show that eQTLs in specific tissues tend to be most enriched among the predicted functional variants in relevant tissues in Roadmap. We further show how these integrated functional scores can be used to derive the most likely cell/tissue type causally implicated for a complex trait using summary statistics from genome-wide association studies, and estimate a tissue-based correlation matrix of various complex traits. We find large enrichment of heritability in functional components of relevant tissues for various complex traits, with FUN-LDA yielding the highest enrichment estimates relative to existing methods. Finally, using experimentally validated functional variants from the literature and variants possibly implicated in disease by previous studies, we rigorously compare FUN-LDA to state-of-the-art functional annotation methods such as GenoSkyline, ChromHMM, Segway, and IDEAS, and show that FUN-LDA has better prediction accuracy and higher resolution compared to these methods. In summary, we describe a new approach and perform rigorous comparisons with the most commonly used functional annotation methods, providing a valuable resource for the community interested in the functional annotation of noncoding variants. Scores for each position in the human genome and for each ENCODE/Roadmap tissue are available from http://www.columbia.edu/~ii2135/funlda.html.

## 1. INTRODUCTION

Understanding the functional consequences of noncoding genetic variation is one of the most important problems in human genetics. Comparative genomics studies suggest that most of the mammalian conserved and recently adapted regions consist of noncoding elements [1, 2, 3]. Furthermore, most of the loci identified in genome-wide association studies fall in noncoding regions and are likely to be involved in gene regulation in a cell type and tissue specific manner [4]. Noncoding variants are also known to play an important role in cancer. Somatic variants in noncoding regions can act as drivers of tumor progression and germline noncoding variants can act as risk alleles [5]. Thus, improved understanding of tissue-specific functional effects of noncoding variants will have implications for multiple diseases and traits.

Prediction of the functional effects of genetic variation is difficult for several reasons. To begin with, there is no single definition of function. As discussed in [6] there are several possible definitions, depending on whether one considers genetic, evolutionary conservation or biochemical perspectives. These different approaches each have limitations and vary substantially with respect to the specific regions of the human genome that they predict to be functional. In particular the genetic approach, based on experimental evaluation of the phenotypic consequence of a sequence alteration (e.g. by measuring the impact of individual alleles on gene expression in a particular context), is low throughput, laborious and may miss elements that lead to phenotypic effects manifest only in rare cells or specific environmental contexts. The evolutionary approach relies on accurate multispecies alignment which makes it challenging to identify certain functional elements, such as distal regulatory elements, although recently several approaches have been developed for primate- or even human-specific elements [7]. An additional limitation of the evolutionary approach is that it is not sensitive to tissue and cell type. Finally, the biochemical approach adopted by projects such as ENCODE [3] and Roadmap Epigenomics [8], although helpful in identifying potentially regulatory elements in specific contexts, does not provide definitive proof of function since the observed biochemical signatures can occur stochastically and in general are not completely correlated with function. Besides the difficulty in precisely defining function, a challenge is that the use of functional genomics features from ENCODE and Roadmap (e.g. ChIP-seq and DNase I hypersensitive sites signals) are mostly useful for predicting the effects of variants in cis-regulatory elements, such as promoters, enhancers, silencers and insulators. Other classes of functional variants, for example those with effects on post-transcriptional regulation by alteration of RNA secondary structure or RNA-protein interactions would be missed by these features.

Recently, several computational approaches have been proposed to predict functional effects of genetic variation in noncoding regions of the genome based on epigenetic and evolutionary conservation features [2, 9, 10, 11]. These predictions are not specific to particular cell types or tissues. Here we are interested in predicting functional effects of genetic variants in specific cell types and tissues. The ENCODE Project and the Roadmap Epigenomics Project have profiled various epigenetic features, including histone modifications and chromatin accessibility, genome-wide in more than a hundred different cell types and tissues. Histone modifications are chemical modifications of the DNA-binding histone proteins that influence transcription as well as other DNA processes. Particular histone modifications have characteristic genomic distributions [12]. For example, trimethylation of histone H3 lysine 4 (H3K4me3) is associated with promoter regions, monomethylation of histone H3 lysine 4 (H3K4me1) is associated with enhancer regions, and acetylation of histone H3 lysine 27 (H3K27ac) and of histone H3 lysine 9 (H3K9ac) are associated with increased activation of enhancer and promoter regions [8]. Repressive marks include H3K27me3 (trimethylation of histone H3 lysine 27) and H3K9me3 (trimethylation of histone H3 lysine 9), both associated with inactive promoters of protein-coding genes; H3K27me3 is found in facultatively repressed genes by Polycomb-group factors, while H3K9me3 is found in heterochromatin regions corresponding to constitutively repressed genes [13]. There are dozens of chromatin marks assayed in large numbers of different cell types and tissues, and studying them individually is inefficient.

Several unsupervised approaches exist for integration of these epigenetic features in specific cell types and tissues. Such integrative approaches reflect the belief that epigenetic features interact with one another to control gene expression. One class of methods attempts to segment the genome into non-overlapping segments, representing major patterns of chromatin marks, and labels these segments using a small set of labels such as active transcription start site, enhancer, strong transcription, weak transcription, quiescent etc. This class includes methods such as ChromHMM [8, 14, 15] and Segway [16], based on Hidden Markov Models (HMMs) and Dynamic Bayesian Networks respectively. ChromHMM is based on complete pooling of data from multiple tissues and fitting a single model to this superdataset, while Segway is based on fitting separate models to data from each tissue (no pooling). Various extensions of these early segmentation approaches have been proposed. Several approaches have focused on better modeling the read count data using Poisson-lognormal and negative multinomial distributions [17, 18], while others have focused on better modeling of the correlations among related cell types and tissues [19, 20, 21]. Yet another approach attempts to improve the HMM parameter estimation procedure in ChromHMM by replacing the EM algorithm with a spectral learning procedure [22]. Another class of methods focuses exclusively on predicting functional effects of variants, rather than segmenting the genome as discussed above. A recent method in this class, GenoSkyline [23], is based on fitting a two-component mixture model of multivariate Bernoulli distributions to epigenetic data for each tissue separately, and then computing a posterior probability for each variant to be in the functional class.

We introduce here a new integrated functional score that combines different epigenetic features in specific cell types and tissues. Our model is based on the latent Dirichlet allocation (LDA) model [24], a generative probabilistic model used often in the topic modeling literature, that allows joint modeling of data from multiple cell types and tissues. The variant scores in each tissue are modeled as a mixture over latent functional classes. In the mixture distribution, we assume that the mixture components are shared across all the tissues, while the mixture proportions for the different functional classes can vary from tissue to tissue (more details on the model and inference algorithm are given in the Methods section). Since our primary goal is to provide a functional score (as opposed to a functional element annotation) we focus on integrating four activating histone modifications (i.e. H3K4me1, H3K4me3, H3K9ac, H3K27ac) and DNase. For the four activating histone modifications data, we compute “valley” scores (Methods), motivated by previous work showing that within regions of high histone acetylation, local minima (or valleys) are strongly associated with transcription factor binding sites [25]. We fit the LDA model with multiple functional classes to these data, and compute for each position its posterior probability to belong to a functional class. We define the functional score at a position as the sum of posterior probabilities for the designated ‘active enhancer’ and ‘active promoter’ classes.

The proposed LDA model has several advantages. First, because the model is fit jointly to data from multiple cell types and tissues, cross-tissue comparisons are meaningful. Second, our method makes no distributional assumptions on the data, allowing us to avoid various data transformations employed by other approaches (such as dichotomization, or other transformations needed to make the data conform more closely to various parametric assumptions), and facilitating the integration of data with arbitrary distributions. Third, by using the valley scores we can improve the precision of locating functional variants relative to methods that utilize smoothed data or peak regions. Fourth, even though we only provide functional scores in the tissues and cell types available in Roadmap, it is easy to perform functional prediction in additional cell types and tissues once the model has been fit to the original Roadmap data. Furthermore, while we regard FUN-LDA as primarily an approach to perform cell type and tissue specific functional prediction in the same sense as the GenoSkyline approach, we explicitly define the functional variants as those falling in ‘active promoter’ or ‘active enhancer’ elements.

In the next section, we demonstrate the usefulness of our predictions using several validation experiments. In summary, we present the following results: (1) we provide cell type and tissue specific functional predictions for every possible position in the hg19 human genome for 127 cell types and tissues in Roadmap, (2) we provide a global view of the sharing of predicted functional variants across large number of cell types and tissues, and show that predicted functional variants that fall in promoters are more likely to be shared across many tissues compared with those that fall in enhancers, (3) we show that eQTLs identified in specific tissues from several sources tend to be most enriched among the predicted functional variants in a relevant Roadmap tissue, (4) we use these cell type and tissue specific scores in conjunction with summary statistics from 21 genome-wide association studies (GWAS) to identify the most likely causal cell type/tissue causally implicated for a particular trait, and estimate a tissue-based correlation matrix among these complex traits, (5) we use experimentally validated functional variants in the literature to rigorously compare FUN-LDA with state-of-the-art functional annotation methods such as GenoSkyline, ChromHMM, Segway, and IDEAS.

## 2. RESULTS

### 2.1. FUN-LDA model with nine classes

Here we use data for four activating histone modifications, namely H3K4me1, H3K4me3, H3K9ac, H3K27ac, and DNase for 127 different cell types and tissues represented in the Roadmap datasets (see Supplemental Tables S1 and S2). Not all of the histone marks were profiled for each of the 127 different cell types and tissues. However, using the relationships between different marks within and across tissues, signal tracks have been predicted for each of these marks across all tissues [8, 15]. We make use of these predicted signal tracks to compute integrated functional scores for every possible position in the human genome for 127 cell types and tissues. Specifically, using the perplexity based criterion (see Methods section) and prior knowledge on the relationship of histone modifications and chromatin states, we investigated models with varying number of classes, and have chosen as our final model a model with nine classes (as shown in Supplemental Figure S1 the perplexity measure begins to plateau starting with models with 9 classes). We fit the LDA model with nine classes to the valley scores for the active histone modification data, and original DNase, and compute posterior probabilities at each position for the different functional classes. The active functional classes correspond to active promoters and active enhancers (Supplemental Figure S2). When comparing with genome segmentation approaches such as ChromHMM (25 state model), Segway and IDEAS we also make a similar partition (see Methods section and Supplemental Table S3). For each position, the sum of the posterior probabilities for the classes in the functional group is used to score the position for both our method and ChromHMM. Segway and IDEAS only provide a functional class assignment for each position for each cell type and tissue in Roadmap, and we use these assignments to identify the functional variants. The proportion of positions in the functional group for each method is shown in Supplemental Figure S3. FUN-LDA, ChromHMM and DNase-narrow (DNase narrow peaks) estimate that an average of 2% of the genome is functional in a cell type or tissue in Roadmap, with the remaining methods producing higher estimates for the size of the functional component.

#### Sharing of predicted functional variants across tissues and cell types

We compute for each variant in the 1000 Genomes project a probability to be in the functional class for each tissue in Roadmap separately. In Figure 1 we provide a global picture of the sharing of predicted functional variants across tissues in Roadmap using the generalized Jaccard similarity index, a measure of overlap between predicted functional variants in two tissues (see Methods section). General tissue groupings are indicated in different colors. As expected, tissues that are functionally related tend to cluster together. There are roughly three major groups: blood cells (indicated in red), including various primary immune cell subtypes, stem cells (indicated in blue) and a third group corresponding to various solid organs (this grouping is also apparent in the multi-dimensional scaling visualization of the correlations between the functional scores in Supplemental Figure S4; see also [8], [14], and Supplemental Figure S5 for related results using single histone marks).

**Figure 1.**
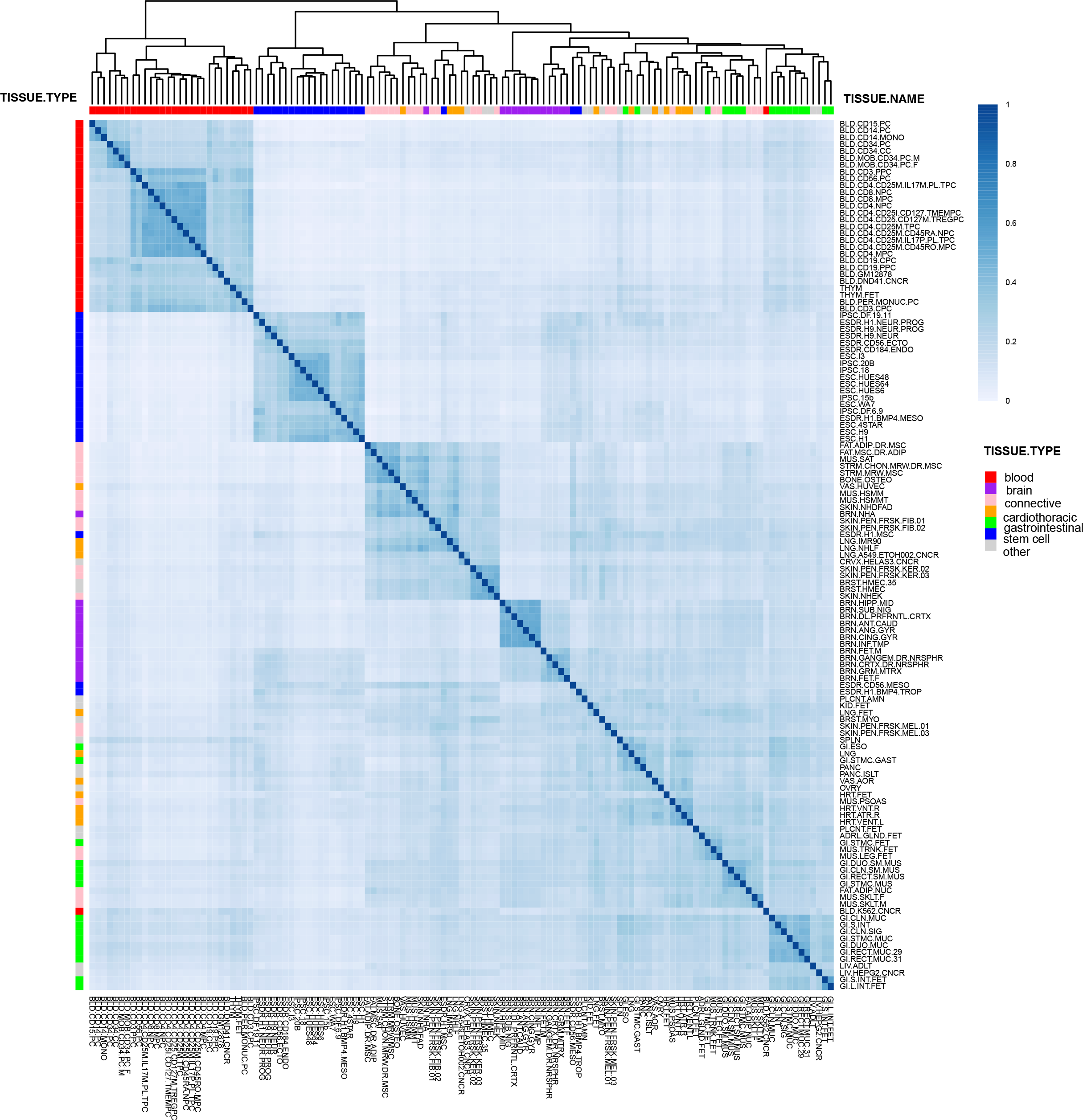
Jaccard index of overlap among functional variants in different cell types and tissues in Roadmap Epigenomics. Hierarchical clustering is used to cluster the different cell types and tissues.

Overall, the median Jaccard index across all pairs of tissues is 0.24. As a comparison, we have also computed the Jaccard overlap indices using predicted functional variants that fall in promoters, and separately in enhancers (Methods section). The median Jaccard index for variants falling in promoters is 0.33, and 0.16 for variants falling in enhancers, concordant with existing literature showing that there tends to be more sharing across tissues for predicted functional variants in promoters vs. those in enhancers [26].

### 2.2. Enrichment analyses using eQTLs from the Genotype-Tissue Expression project, Geuvadis and TwinsUK data.

#### eQTLs from the Genotype-Tissue Expression project

The Genotype-Tissue Expression (GTEx) project is designed to establish a comprehensive data resource on genetic variation, gene expression and other molecular phenotypes across multiple human tissues [27]. We focus here on the cis-eQTL results from the GTEx V6 release comprising RNA-seq data on 7,051 samples in 44 tissues, each with at least 70 samples (Supplemental Table S4). We are interested in identifying for each GTEx tissue the Roadmap tissue that is most enriched in eQTLs from that GTEx tissue relative to other Roadmap tissues (see Methods section). We exclude from analysis the sex-specific GTEx tissues, most of which have no relevant counterpart in Roadmap. These include the following tissues: ovary, vagina, uterus, testis, prostate, breast. In Table 1 we show the top Roadmap tissue for each remaining GTEx tissue, along with the p value from the enrichment test. In most cases, eQTLs from a GTEx tissue show the most enrichment in the functional component of a relevant Roadmap tissue. For example, for liver tissue in GTEx, liver is the Roadmap tissue with the highest enrichment, for pancreas tissue in GTEx, the Roadmap tissue with the highest enrichment is pancreas, for skeletal muscle tissue in GTEx, the most enriched Roadmap tissue is skeletal muscle. However, there are also a few cases where the top tissue is not necessarily the most intuitive one, such is the case for lung and several brain tissues. Generally, the tissues with unexpected combinations tend to either have small sample sizes for eQTL discovery in GTEx (such as brain tissues) or inadequate representation in Roadmap (e.g. thyroid, pituitary gland, artery - tibial, artery - coronary, esophagus - gastroesophageal junction etc.). Most of the mismatches have relatively large p values as well *(p >* 0.001).

**Table 1:** Enrichment of eQTLs from different sources (GTEx, Geuvadis and TwinsUK cohort) among FUN-LDA predicted functional variants in tissues and cell types in Roadmap Epigenomics. The top Roadmap tissue is given for each eQTL tissue, along with the p value from a two-sample proportion test.

| Study | Tissue | Roadmap Epigenome Name | -log<sub>10</sub>(p) |
| --- | --- | --- | --- |
| GTEx | Whole Blood | Primary neutrophils from peripheral blood | 189.72 |
|  | Cells - Transformed fibroblasts | Muscle Satellite Cultured Cells | 62.69 |
|  | Cells - EBV-transformed lymphocytes | GM12878 Lymphoblastoid Cells | 37.74 |
|  | Liver | Liver | 31.82 |
|  | Muscle - Skeletal | Skeletal Muscle Male | 19.42 |
|  | Heart - Left Ventricle | Fetal Heart | 15.83 |
|  | Esophagus - Mucosa | Esophagus | 12.78 |
|  | Pancreas | Pancreas | 10.84 |
|  | Colon - Transverse | Rectal Mucosa Donor 31 | 10.46 |
|  | Artery - Tibial | Stomach Smooth Muscle | 7.74 |
|  | Esophagus Muscularis | Stomach Smooth Muscle | 6.74 |
|  | Thyroid | Fetal Intestine Small | 5.96 |
|  | Skin - Sun Exposed (Lower leg) | Foreskin Keratinocyte Primary Cells skin03 | 5.47 |
|  | Spleen | Primary B cells from peripheral blood | 5.35 |
|  | Artery - Aorta | Aorta | 5.28 |
|  | Brain - Hippocampus | Brain Cingulate Gyrus | 5.10 |
|  | Small Intestine - Terminal Ileum | Fetal Intestine Large | 5.04 |
|  | Heart - Atrial Appendage | Fetal Heart | 4.90 |
|  | Adipose - Subcutaneous | Adipose Nuclei | 4.74 |
|  | Colon - Sigmoid | Colon Smooth Muscle | 4.62 |
|  | Brain - Caudate (basal ganglia) | Brain Substantia Nigra | 4.17 |
|  | Brain - Cerebellum | Adipose Derived Mesenchymal Stem Cell Cultured Cells | 4.12 |
|  | Nerve - Tibial | Brain Hippocampus Middle | 4.11 |
|  | Adrenal Gland | Fetal Adrenal Gland | 3.94 |
|  | Skin - Not Sun Exposed (Suprapubic) | Foreskin Keratinocyte Primary Cells skin03 | 3.56 |
|  | Brain - Putamen (basal ganglia) | Brain Substantia Nigra | 3.36 |
|  | Brain - Cerebellar Hemisphere | Brain Angular Gyrus | 3.08 |
|  | Stomach | Stomach Mucosa | 3.02 |
|  | Lung | Osteoblast Primary Cells | 2.57 |
|  | Brain - Cortex | Mesenchymal Stem Cell Derived Chondrocyte Cultured Cells | 2.10 |
|  | Adipose - Visceral (Omentum) | Primary T helper cells from peripheral blood | 2.00 |
|  | Pituitary | Primary T helper cells PMA-I stimulated | 1.96 |
|  | Brain - Nucleus accumbens (basal ganglia) | H9 Cells | 1.80 |
|  | Esophagus - Gastroesophageal Junction | Primary neutrophils from peripheral blood | 1.64 |
|  | Brain - Frontal Cortex (BA9) | NHDF-Ad Adult Dermal Fibroblast Primary Cells | 1.61 |
|  | Artery - Coronary | Primary B cells from peripheral blood | 1.35 |
|  | Brain - Hypothalamus | Osteoblast Primary Cells | 1.29 |
|  | Brain - Anterior cingulate cortex (BA24) | A549 EtOH 0.02pct Lung Carcinoma Cell Line | 1.04 |
| Geuvadis | Lymphoblastoid cell line | GM12878 Lymphoblastoid Cells | 8.57 |
| TwinsUK | Blood | Primary neutrophils from peripheral blood | 7.54 |
|  | Fat | Mesenchymal Stem Cell Derived Adipocyte Cultured Cells | 6.80 |
|  | Skin | Foreskin Keratinocyte Primary Cells skin02 | 3.62 |
|  | Lymphoblastoid cell line | GM12878 Lymphoblastoid Cells | 3.08 |

#### eQTLs from the Geuvadis and TwinsUK data

We sought to perform similar analyses using eQTLs identified in other studies, in particular in lymphoblastoid cell lines (LCLs) in the Geuvadis project, and four tissues (fat, lymphoblastoid cell lines, skin and whole blood) using individuals from the TwinsUK cohort. We have focused here on the lead eQTLs (those variants most associated with expression levels [28]), and performed similar enrichment analyses as for the eQTLs from GTEx. As shown in Table 1, the most enriched Roadmap tissue corresponds very well to the tissue of origin used in the eQTL discovery, providing an independent validation of the findings using the eQTLs from GTEx.

### 2.3. Prediction of causal tissues for 21 complex traits

As an application of our scores to complex trait genetics, we use the recently developed stratified linkage disequilibrium (LD) score regression framework [29] to identify the most relevant cell types and tissues for 21 complex traits for which moderate to large GWAS studies have been performed (Table 2; [30]-[50]). The stratified LD score regression approach uses information from all single nucleotide polymorphisms (SNPs) and explicitly models LD to estimate the contribution to heritability of different functional classes of variants. We modify this method to weight SNPs by their tissue specific functional score (e.g. FUN-LDA), and in this way we assess the contribution to heritability of predicted functional SNPs in a particular Roadmap cell type or tissue (see Supplemental Material for more details).

**Table 2:**
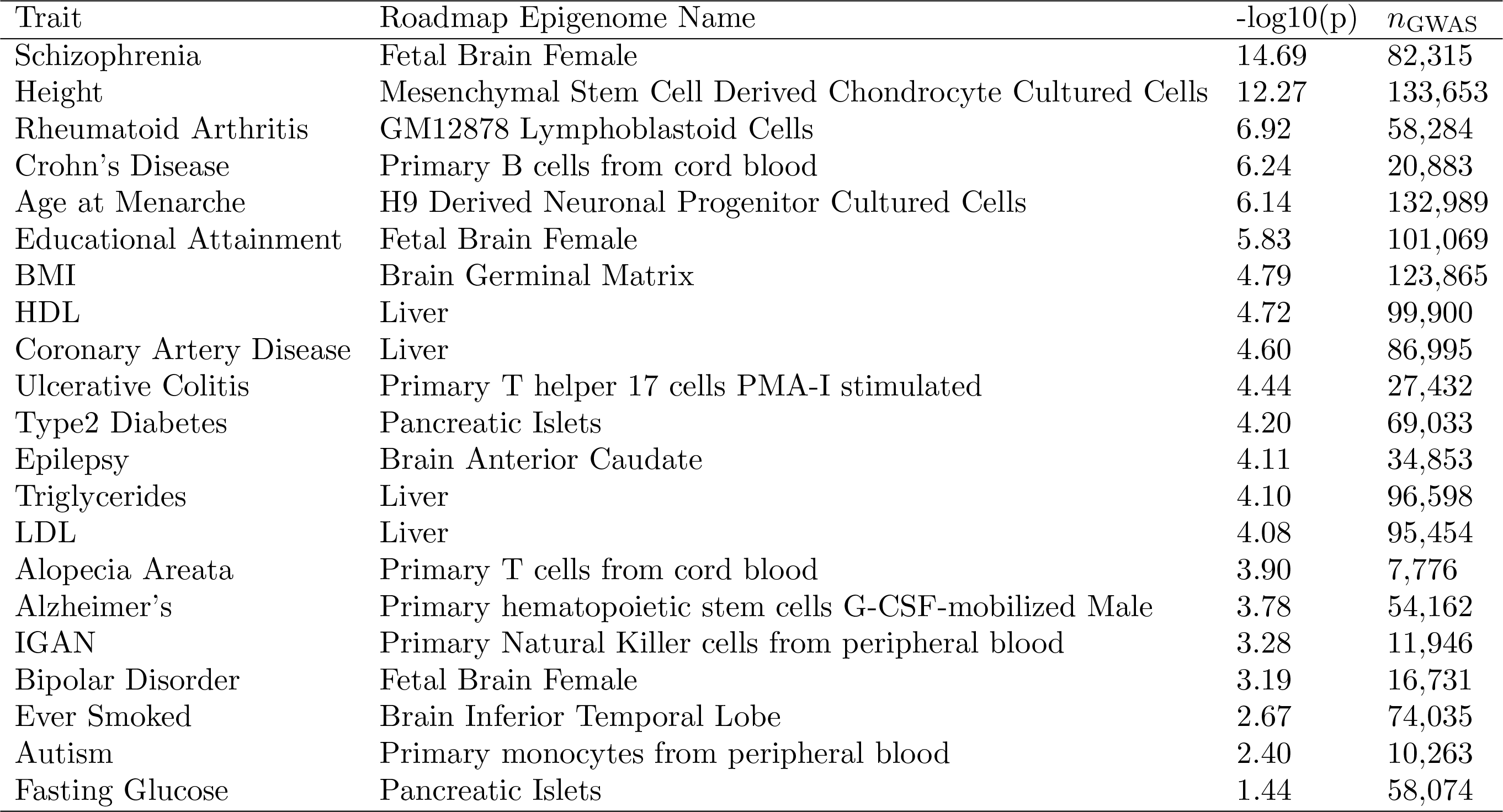
Top cell type/tissue in Roadmap for 21 GWAS traits using FUN-LDA posterior probabilities. The p value from the stratified LD score regression, as well as the GWAS sample size are reported for each trait.

| Trait | Roadmap Epigenome Name | $-\log_{10}(p)$ | $n_{\text{GWAS}}$ |
| --- | --- | --- | --- |
| Schizophrenia | Fetal Brain Female | 14.69 | 82,315 |
| Height | Mesenchymal Stem Cell Derived Chondrocyte Cultured Cells | 12.27 | 133,653 |
| Rheumatoid Arthritis | GM12878 Lymphoblastoid Cells | 6.92 | 58,284 |
| Crohn's Disease | Primary B cells from cord blood | 6.24 | 20,883 |
| Age at Menarche | H9 Derived Neuronal Progenitor Cultured Cells | 6.14 | 132,989 |
| Educational Attainment | Fetal Brain Female | 5.83 | 101,069 |
| BMI | Brain Germinal Matrix | 4.79 | 123,865 |
| HDL | Liver | 4.72 | 99,900 |
| Coronary Artery Disease | Liver | 4.60 | 86,995 |
| Ulcerative Colitis | Primary T helper 17 cells PMA-I stimulated | 4.44 | 27,432 |
| Type2 Diabetes | Pancreatic Islets | 4.20 | 69,033 |
| Epilepsy | Brain Anterior Caudate | 4.11 | 34,853 |
| Triglycerides | Liver | 4.10 | 96,598 |
| LDL | Liver | 4.08 | 95,454 |
| Alopecia Areata | Primary T cells from cord blood | 3.90 | 7,776 |
| Alzheimer's | Primary hematopoietic stem cells G-CSF-mobilized Male | 3.78 | 54,162 |
| IGAN | Primary Natural Killer cells from peripheral blood | 3.28 | 11,946 |
| Bipolar Disorder | Fetal Brain Female | 3.19 | 16,731 |
| Ever Smoked | Brain Inferior Temporal Lobe | 2.67 | 74,035 |
| Autism | Primary monocytes from peripheral blood | 2.40 | 10,263 |
| Fasting Glucose | Pancreatic Islets | 1.44 | 58,074 |

In Table 2 we show the top Roadmap cell type/tissue (the one with the smallest p value from testing whether predicted functional variants in a tissue contribute significantly to SNP heritability) for each of the 21 complex traits using FUN-LDA to predict functional variants in specific cell types and tissues. For most disorders, the top tissue has previously been implicated in their pathogenesis. For example, the top tissues for body mass index (BMI) are brain tissues, consistent with recent findings indicating that BMI-associated loci are enriched for expression in the brain and central nervous system [51]. Similarly, brain represents the top tissue for most neuropsychiatric disorders, education levels, and smoking. Blood-derived and immune cells represent the top tissue for virtually all of the autoimmune conditions available for analysis. For example, GWAS findings for ulcerative colitis map specifically to the regulatory elements in Th17 cells, whereas lymphoblastoid cell lines represent the top cell type for rheumatoid arthritis. Another interesting finding involves primary hematopoietic stem cells for Alzheimer’s disease, consistent with emerging data on the involvement of bone marrow-derived immune cells in the pathogenesis of neurodegeneration [53].

Results for other methods are shown in Supplemental Tables S5-S7. Estimates of enrichment (defined as the proportion of SNP heritability in the category divided by the proportion of SNPs in that category) for the functional component in the top tissues in Supplemental Tables S5-S7 are shown in Figure 2. On average across traits, the functional component for the top tissue as defined by FUN-LDA shows the highest enrichment relative to other methods, with approximately 2% of the SNPs (functional in the top tissue) explaining an estimated 32% of SNP heritability. FUN-LDA is followed closely by DNase-narrow and ChromHMM. Methods such as DNase-gapped, GenoSkyline and IDEAS show substantially lower enrichments; e.g. for IDEAS, 7.1% SNPs explain an estimated 52% of heritability. In terms of the top tissues identified by each method, it is difficult to make an objective comparison since the underlying tissues and cell types are not known for many complex traits. However looking at the results in Supplemental Tables S5-S7, one can point out several likely mismatches, such as ‘Lung’ for coronary artery disease identified by both GenoSkyline and DNase-narrow, or ‘Dnd41 T-Cell Leukemia Cell Line’ and ‘Fetal Thymus’ identified for epilepsy by DNase and DNase-narrow, respectively. Notably, for Type 2 Diabetes, FUN-LDA, Segway and DNase-gapped were the only methods to point to pancreatic tissue.

**Figure 2.**
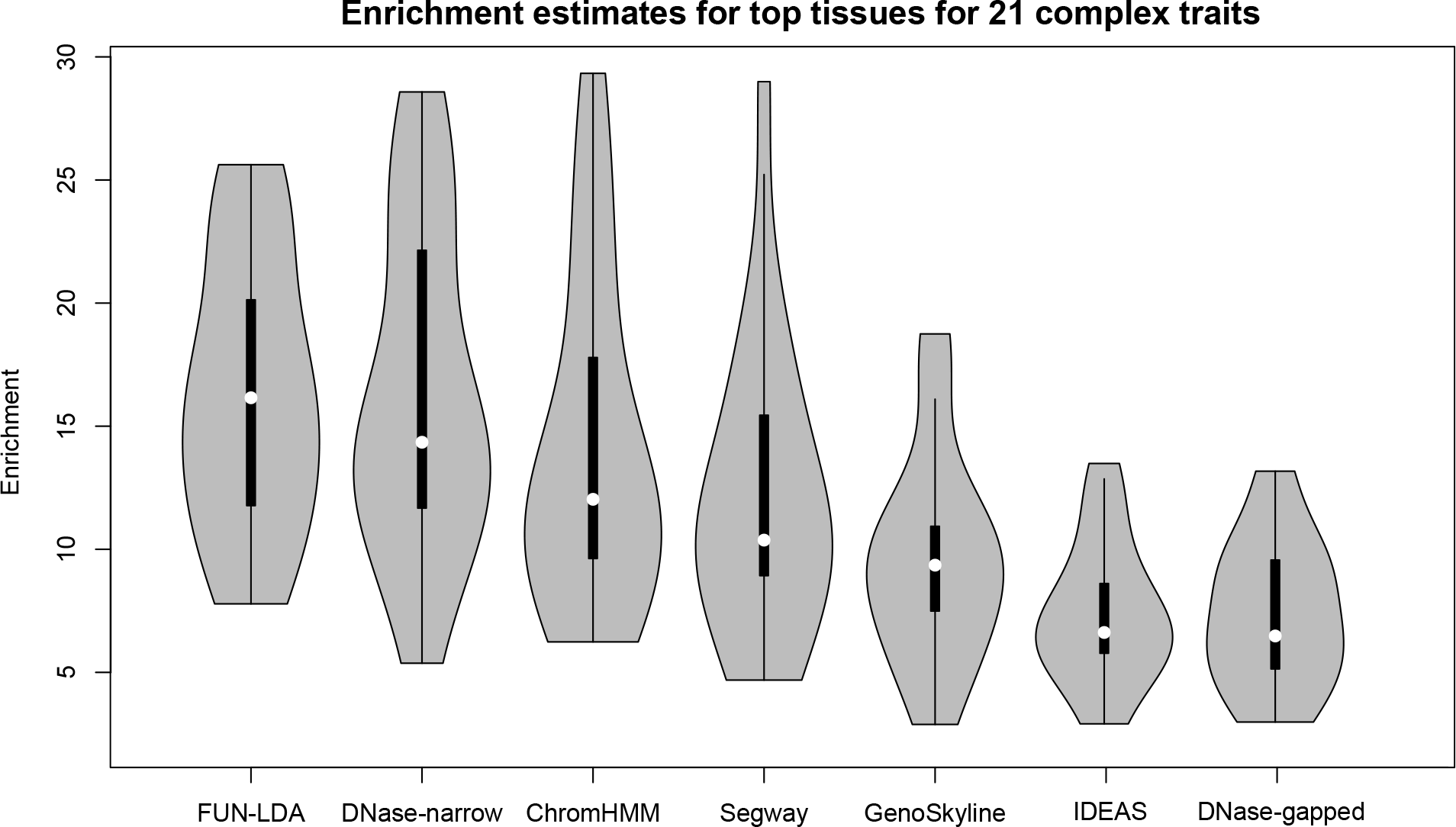
Enrichment estimates (the proportion of SNP heritability in the functional component divided by the proportion of SNPs in that component) for different methods across top tissues for 21 complex traits. Enrichment estimates for DNase are omitted since they do not make sense for continuous annotations, such as DNase.

In Figure 3 we show the correlation matrix for the 21 traits based on the Z-scores from the LD score regressions (see Methods section for more details on how these pairwise correlations were estimated). This correlation matrix reflects the extent to which traits share the same causal tissues, rather than the genetic correlation [54]. Three large phenotypic clusters are clearly evident. The most tightly correlated cluster contains autoimmune and inflammatory conditions, including Crohn’s disease, alopecia areata, rheumatoid arthritis and IgA nephropathy. As expected, these conditions share highest functional scores in blood-derived immune cells. The second most strongly inter-correlated cluster is driven by scores in neuronal tissues, and consists of BMI, age at menarche, educational attainment, schizophrenia, and smoking history, with somewhat weaker correlations with autism, epilepsy and bipolar disorder. Lastly, there is a clear co-clustering of cardio-metabolic traits that map to the tissues of liver, pancreas, and small intestine. Also, as shown, Alzheimer’s disease clusters with LDL, HDL and triglycerides, concordant with recent reports on a link between cardio-vascular disease and Alzheimer’s disease [55].

**Figure 3.**
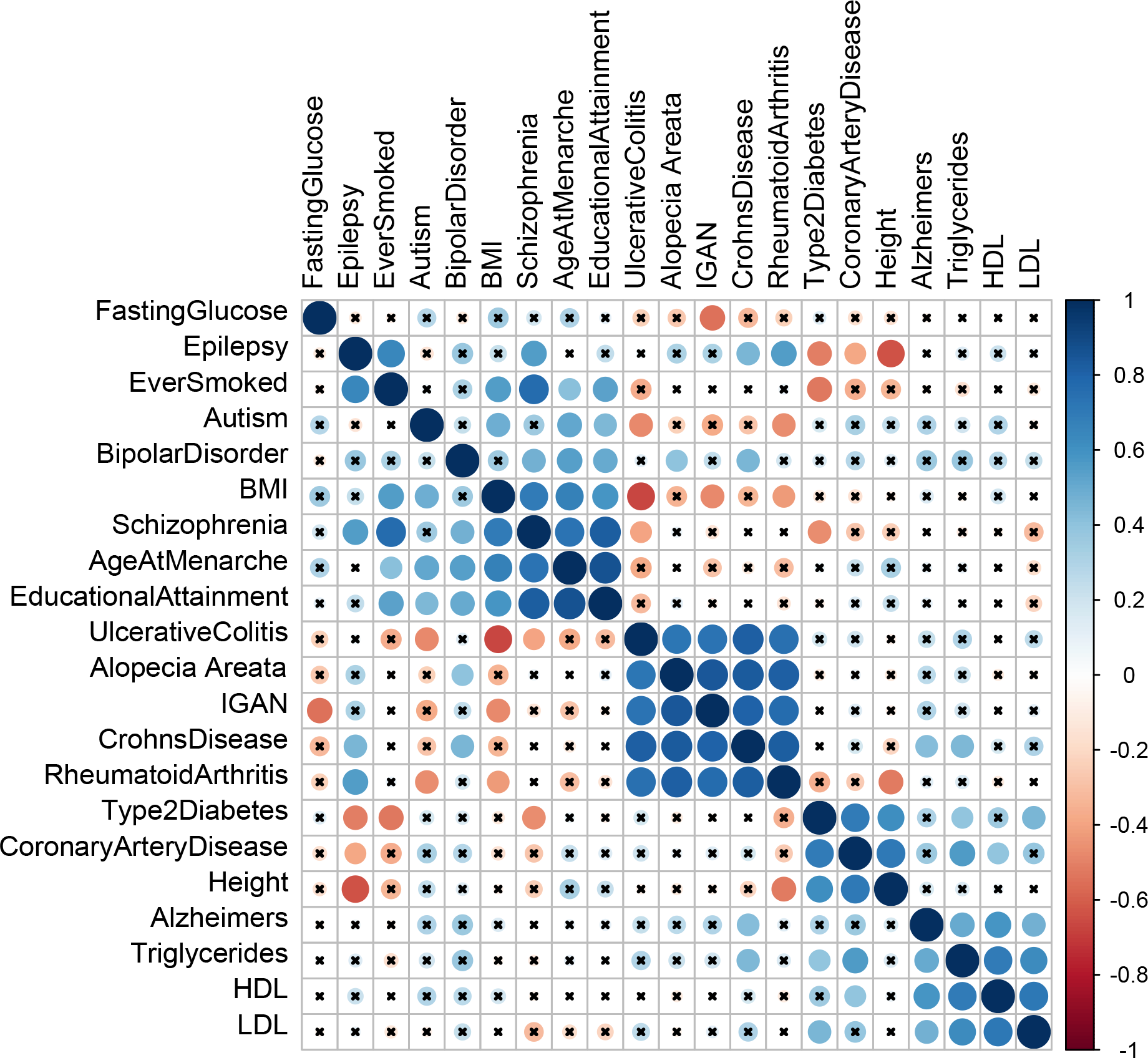
Tissue Correlations for 21 common traits. Hierarchical clustering (average linkage method) is used to cluster diseases. The ‘x’ symbol indicates that those correlations are not significant at the 0.0001 level.

### 2.4. Validations of our model’s predictions, and comparisons with existing methods for functional annotation

To further assess the accuracy of our predictions and compare with existing approaches we use variants in the literature that have been experimentally shown to have a regulatory function. We focus on several main lists of variants: (a) eight variants implicated in Mendelian and complex diseases, with additional experimental validation of their functional effects [56]-[63], (b) confirmed regulatory variants from a multiplexed reporter assay in lymphoblastoid cell lines [64], (c) regulatory motifs in 2, 000 predicted human enhancers using a massively parallel reporter assay in two human cell lines, liver carcinoma (HepG2) and erythrocytic leukemia (K562) cell lines [65], and (d) validated enhancers in 167 ultra conserved sequence elements [66].

#### 2.4.1. Noncoding variants implicated in Mendelian and complex traits with experimentally predicted regulatory function

We selected the following eight SNPs that have been shown experimentally to have a regulatory function in particular tissues: rs6801957 [56], rs12821256 [57], rs12350739 [58], rs12740374 [59], rs356168 [60], rs2473307 [61], rs227727 [62], and rs144361550 [63]. In Figure 4, Supplemental Figures S6-S11 we show the predictions in ~ 2 kb windows centered at these SNPs from the different approaches: FUN-LDA, GenoSkyline, ChromHMM (25 state model), Segway and IDEAS. For each of these SNPs, we select the tissue in Roadmap that we believe is closest to the tissue used in the original functional studies ([56]-[63], Supplemental Table S8). We summarize below the results for two of the SNPs, rs6801957 and rs12821256, that show more tissue specificity relative to the other SNPs in the set (i.e. are predicted to be functional in a small number of Roadmap tissues). For the remaining six SNPs the results are summarized in the Supplemental Material, and Supplemental Figures S6-S11.

**Figure 4.**
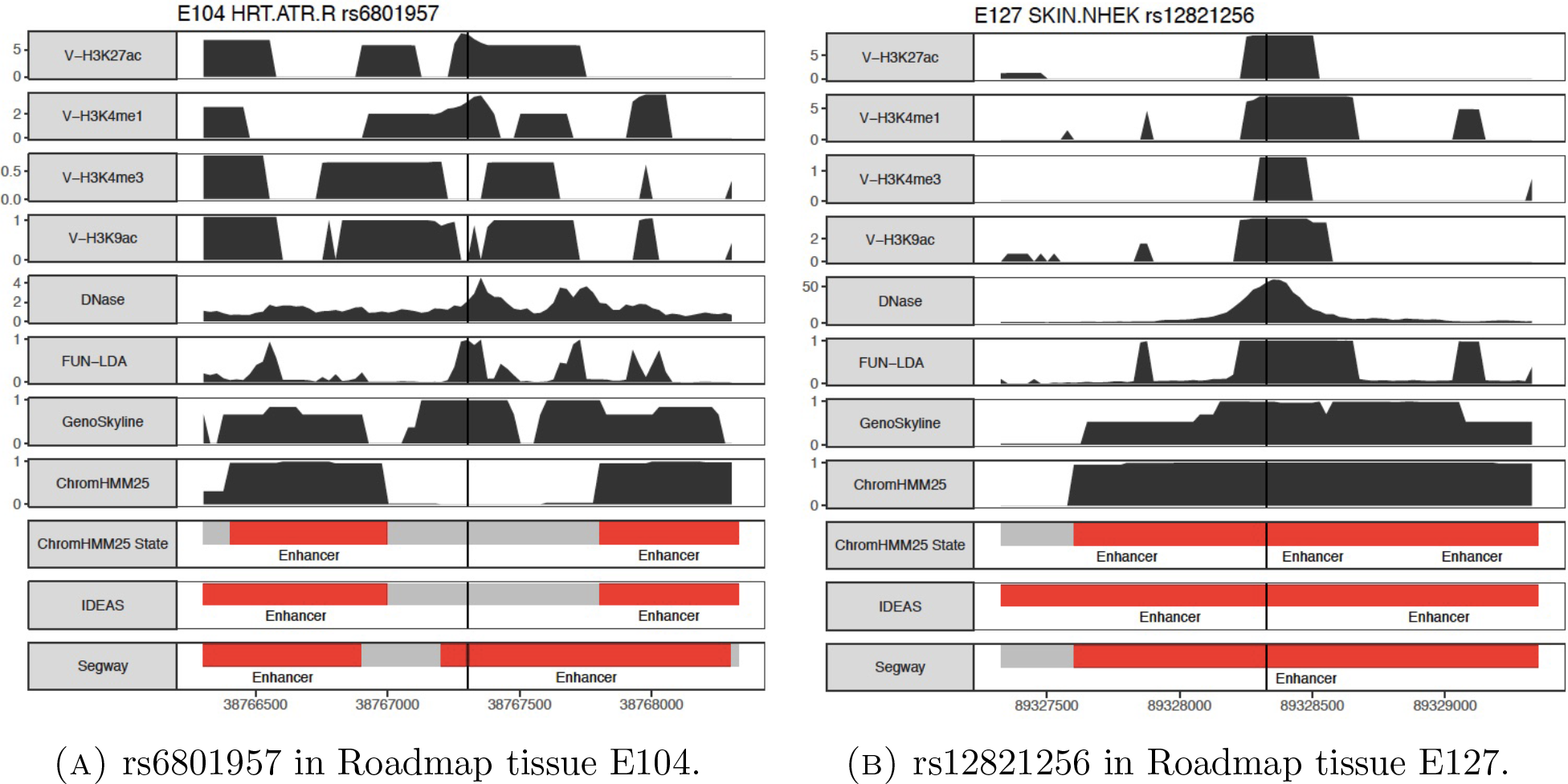
Valley scores for four activating histone marks and DNase, posterior probabilities from FUN-LDA, GenoSkyline, and ChromHMM (25 state model), and segmentations from ChromHMM, IDEAS and Segway are shown in 2 kb windows centered around the lead SNPs. For clarity we only highlight in the segmentations the type of states we consider functional (enhancer states in red, promoter states in blue) for the different segmentation approaches.

rs6801957: In [56], the authors show that this SNP, found associated in GWAS studies with ECG measures, is associated with lower *SCN5A* expression in heart tissue in humans and mice. In Figure 4 we show the predictions for Roadmap tissue E104, Right Atrium.

rs12821256, a SNP associated with blond hair color in Iceland and the Netherlands, is located in an enhancer and influences expression of the *KITLG* gene in cultured human keratinocytes [57]. In Figure 4 we show the predictions for Roadmap tissue E127, NHEK-Epidermal Keratinocyte Primary Cells.

For both SNPs, FUN-LDA assigns a posterior probability of 1 to be functional in the corresponding tissues. Compared with existing methods, the region predicted functional by FUN-LDA tends to be substantially smaller, and therefore FUN-LDA has better ability to predict the causal variant in a region of interest relative to existing approaches.

#### 2.4.2. Confirmed regulatory variants (emVars) from a multiplexed reporter assay

In [64], the authors have applied a new version of the massively parallel reporter assay (MPRA) to identify variants with effects on gene expression. In particular, they apply it to 32, 373 variants from 3, 642 cis-expression quantitative trait loci and control regions in lymphoblastoid cell lines (LCLs), and identify 842 variants showing differential expression between alleles, or emVars, expression-modulating variants. We use this set of 842 emVars as positive control variants. Our negative control variants are those variants tested using the MPRA where neither allele showed differential expression relative to the control, applying a threshold of 0.1 for the Bonferroni corrected p value. After removing from the list of positive and negative control variants those variants that we could not map to a genomic location using the Ensembl database (http://grch37.ensembl.org/index.html), there remained 693 positive control variants and 22, 384 negative control variants.

We compute AUC values for several methods, including FUN-LDA, GenoSkyline, ChromHMM (25 state model), Segway and IDEAS. For ChromHMM we partition the twenty-five states into two groups, ‘functional’ and ‘non-functional’, with the functional group consisting of ‘TssA’, ‘PromU’, ‘PromD1’, ‘PromD2’, ‘EnhA1’, ‘EnhA2’, ‘EnhAF’. For each variant, the sum of ChromHMM posterior probabilities for the classes in the functional group above is used to score the variant. For FUN-LDA we similarly group the designated ‘active promoters’ and ‘active enhancers’ classes to form the ‘functional’ class (see Methods section and Supplemental Table S3). Segway and IDEAS only provide a functional class assignment for each position, and we use these assignments to identify the functional variants. Results are shown in Table 3. As shown, FUN-LDA has higher AUC compared to the existing methods, ChromHMM, GenoSkyline, IDEAS and Segway. Compared with DNase, FUN-LDA performs significantly better than the two binarized versions DNase-narrow and DNase-gapped, the two versions normally used in practice, but it does not outperform the original DNase (on the -log10(p value) scale).

**Table 3:** AUC values for discriminating between variants likely to be functional and control variants. Results are shown for several datasets (three tissues) with experimental validation (MPRA) of potential regulatory variants. Methods include FUN-LDA, GenoSkyline, ChromHMM (25 state model), Segway, IDEAS, and DNase (original, -narrow and -gapped).

| Dataset | Method | AUC |
| --- | --- | --- |
| emVars in [64], E116 | FUN-LDA | 0.709 |
|  | GenoSkyline | 0.662 |
|  | ChromHMM | 0.668 |
|  | Segway | 0.624 |
|  | IDEAS | 0.621 |
|  | DNase | 0.716 |
|  | DNase-narrow | 0.629 |
|  | DNase-gapped | 0.653 |
| Regulatory motifs in [65], E118/HepG2 | FUN-LDA | 0.694 |
|  | GenoSkyline | 0.629 |
|  | ChromHMM | 0.608 |
|  | Segway | 0.618 |
|  | IDEAS | 0.546 |
|  | DNase | 0.719 |
|  | DNase-narrow | 0.561 |
|  | DNase-gapped | 0.550 |
| Regulatory motifs in [65], E123/K562 | FUN-LDA | 0.646 |
|  | GenoSkyline | 0.620 |
|  | ChromHMM | 0.634 |
|  | Segway | 0.585 |
|  | IDEAS | 0.615 |
|  | DNase | 0.654 |
|  | DNase-narrow | 0.524 |
|  | DNase-gapped | 0.565 |

#### 2.4.3. Regulatory motifs in 2,000 predicted human enhancers using a massively parallel reporter assay

In [65], the authors use a massively parallel reporter assay to measure the transcriptional levels produced by targeted motif disruptions in 2,104 candidate enhancers in two human cell lines, liver carcinoma (HepG2) and erythrocytic leukemia (K562) cell lines, providing one of the largest resource of experimentally validated enhancer manipulations in human cells. We use as positive control variants those variants where the p value comparing expression values for the sequence with the motif compared to sequences with scrambled versions of the motif was less than 0.05. We use as negative control variants those variants where this p value was greater than 0.1. After removing those variants whose genomic coordinates we could not resolve, there remained, for HepG2, 525 positive and 1, 451 negative control variants, and for K562, 342 positive and 1, 578 negative control variants. For all methods, we calculate the scores for these motifs by averaging across all bases in the motifs. As shown in Table 3, FUN-LDA has better accuracy compared with GenoSkyline, ChromHMM, IDEAS and Segway, and for HepG2 the improvement is substantial.

We have attempted to form the functional group in an objective manner, based on prior knowledge on what functional classes from the different segmentation approaches (ChromHMM, Segway and IDEAS) should be considered active functional elements. We have performed an additional analysis where we have computed the AUC for all combinations of states (with individual AUC ≥ 0.5) for each segmentation method and selected the state combination with highest AUC for the three datasets above. The results from these analyses are shown in Supplemental Table S10. As shown, even with this optimized state combination, the AUCs for the various methods is most of the times less than for our (unbiasedly selected) state combination for FUN-LDA. Furthermore, the state combination with the maximum AUC often contains states like poised/bivalent promoters that would not a priori be considered functional.

#### 2.4.4. Ultra conserved sequence elements

In [66], the authors used extreme evolutionary sequence conservation as a filter to identify putative gene regulatory sequences. Using this approach, they identified 167 ultra conserved sequence elements, and then used transgenic mouse enhancer assay that links each of these candidate elements to a mouse promoter fused to a lacZ reporter gene. In total, 75 out of 167 candidate sequences functioned reproducibly as tissue-specific enhancers of gene expression by the read out of lacZ expression at mouse embryonic day 11.5. Out of 75 positive fragments, 50 mapped to a single anatomical structure in the E11.5 embryonic tissue, while the remaining 25 enhancers directed expression to two or more anatomical structures. Here, we compare the functional scores for the variants falling into these 75 positive enhancers with scores of variants in the remaining 92 elements. In Table 4 we show the top Roadmap tissue for each method and the corresponding AUC values. Notably, most methods, including FUN-LDA, select embryonic tissue as the top tissue, consistent with the conducted experiment. Importantly, FUN-LDA outperforms all other methods except for GenoSkyline in predicting functional elements based on these enhancer assays.

**Table 4:**
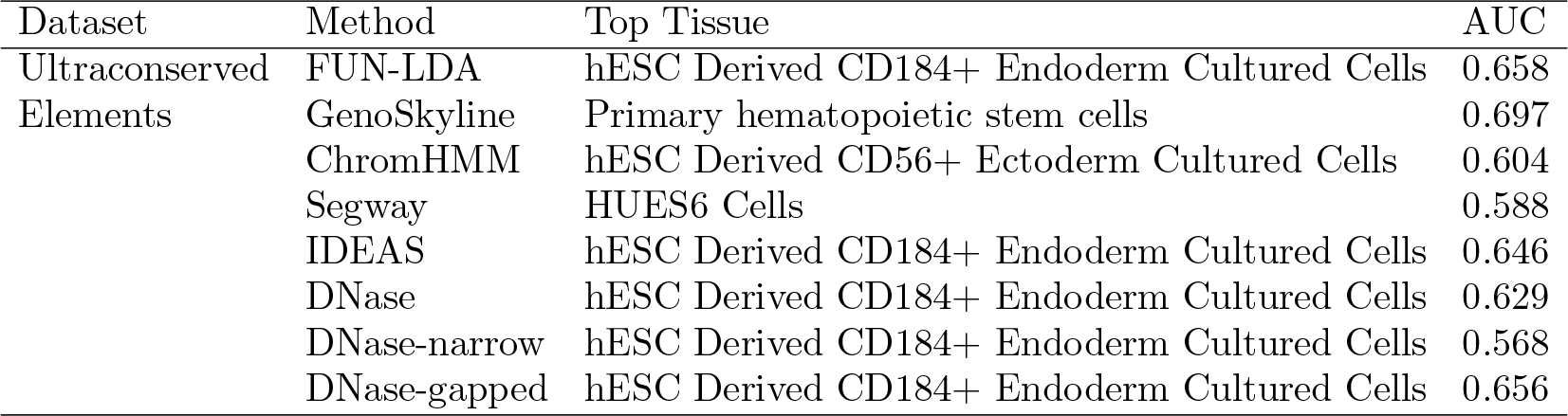
AUC values for discriminating between variants likely to be functional and control variants. Results are shown for validated enhancers in ultra conserved sequence elements [66]. Methods include FUN-LDA, GenoSkyline, ChromHMM (25 state model), Segway, IDEAS, and DNase (original, -narrow and -gapped). The tissue with the highest AUC for each method is also shown.

| Dataset | Method | Top Tissue | AUC |
| --- | --- | --- | --- |
| Ultraconserved<br>Elements | FUN-LDA | hESC Derived CD184+ Endoderm Cultured Cells | 0.658 |
|  | GenoSkyline | Primary hematopoietic stem cells | 0.697 |
|  | ChromHMM | hESC Derived CD56+ Ectoderm Cultured Cells | 0.604 |
|  | Segway | HUES6 Cells | 0.588 |
|  | IDEAS | hESC Derived CD184+ Endoderm Cultured Cells | 0.646 |
|  | DNase | hESC Derived CD184+ Endoderm Cultured Cells | 0.629 |
|  | DNase-narrow | hESC Derived CD184+ Endoderm Cultured Cells | 0.568 |
|  | DNase-gapped | hESC Derived CD184+ Endoderm Cultured Cells | 0.656 |

#### 2.4.5. Widths of predicted functional regions for each method

In Figure 5 we show the distribution of the widths of predicted functional regions including validated functional variants from the three lists above. The width of the functional region around a variant was determined by finding the width of the window around the variant in which the value of the score is greater than 0.5. Widths are truncated at 20, 000 base pairs (so all widths greater than 20, 000 base pairs are represented as 20, 000 base pairs). The FUN-LDA predicted regions are predicted to be substantially narrower compared to the other methods, hence FUN-LDA has the ability to more precisely and more accurately identify the functional variants in a region of interest compared with existing methods.

**Figure 5.**
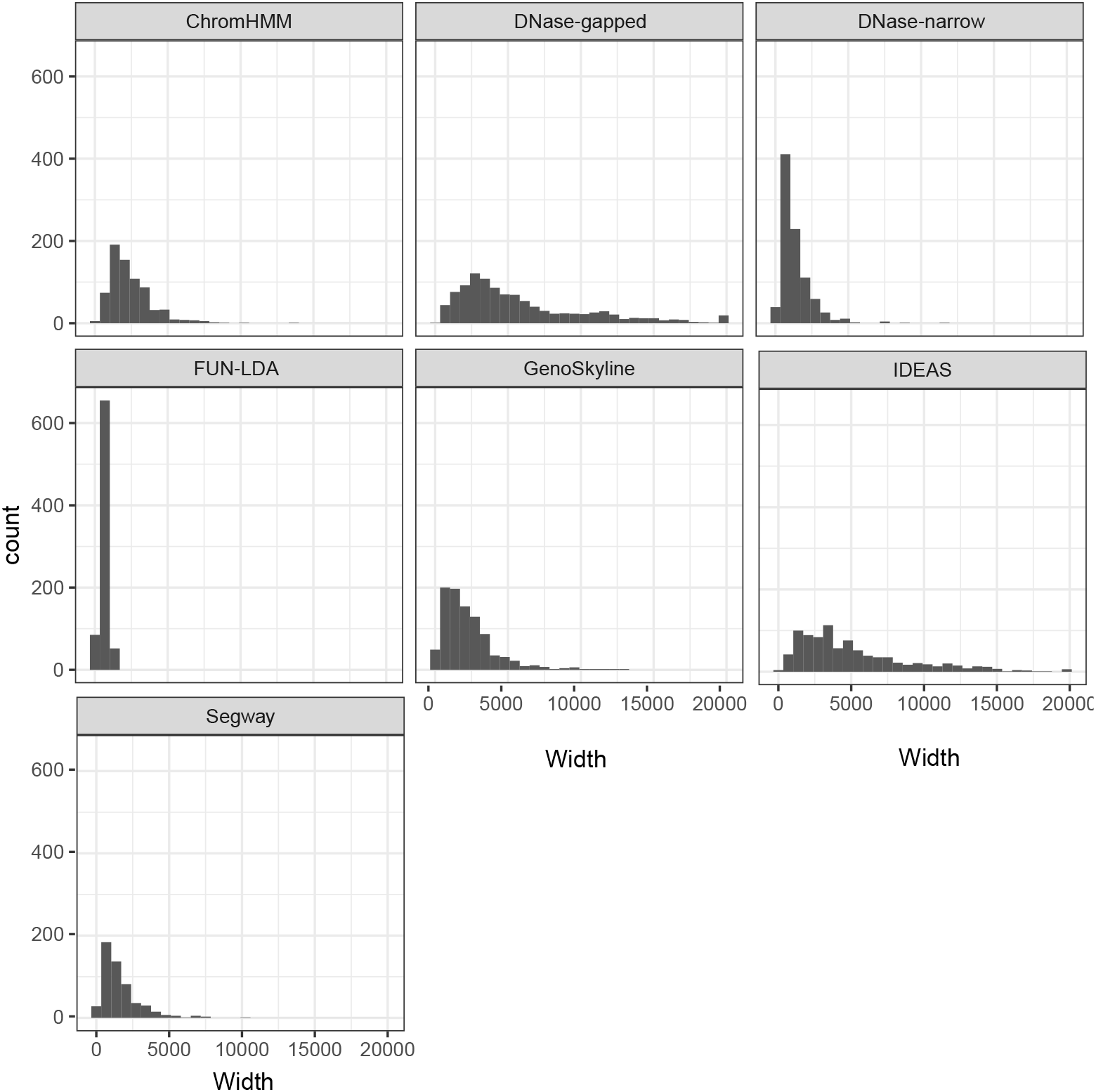
Widths of predicted functional regions (in bps) including validated functional variants from [64], [65] and the eight confirmed variants in Supplemental Table S8.

## 3. DISCUSSION

We have introduced here a new unsupervised approach FUN-LDA for the functional prediction of genetic variation in specific cell types and tissues using histone modification and DNase data from the ENCODE and Roadmap Epigenomics projects, and have provided comparisons with commonly used functional annotation methods. FUN-LDA is based on a mixture model that focuses on identifying the narrow regions in the genome whose disruption is most likely to interfere with function in a particular cell type or tissue. Such context specific functional prediction of genetic variation is essential for understanding the function of noncoding variation across cell types and tissues, and for the interpretation of genetic variants uncovered in GWAS and sequencing studies. While existing segmentation approaches can be used to derive a numeric functional score as well, we have shown that they tend to be less accurate at predicting functional effects, and tend to predict wider functional regions compared to the proposed approach. Relative to other recently developed functional scores, such as GenoSkyline, FUN-LDA has substantially better prediction accuracy, and furthermore makes explicit which classes are considered functionally active, namely active promoters and active enhancers, providing an attractive tool for functional scoring of variants.

In terms of prediction accuracy, we have shown FUN-LDA to outperform existing methods, sometimes substantially. We have also shown that DNase can have higher predictive power than FUN-LDA with respect to the MPRA experiments. However not being a probabilistic score is a significant deficiency of DNase (e.g. enrichment analyses as shown here for eQTL and LD score regression are more difficult to implement/interpret) and in practice researchers are normally using DNase peaks rather than the raw DNase scores, and our method significantly outperforms DNase peaks on the metrics we considered. We note also that the experimental datasets we use here (from the MPRA experiments, and validated enhancers in ultra conserved regions) do not have gold standard labels (for example, the sensitivity for the MPRA assay in [64] is estimated to be between 9% – 24%), and therefore there is an upper limit to the AUC we can achieve on these datasets even with an optimal method.

These cell type and tissue specific functional scores have numerous applications. We have shown here for the first time in the literature that eQTLs from several large studies such as GTEx, Geu-vadis and TwinUK cohort, are most enriched in the functional components from relevant Roadmap tissues. As shown before in [29], and as illustrated here as well, they can be used to infer the most relevant cell types and tissues for a trait of interest, and can help focus the search for causal variants in complex traits by restricting the set of candidate variants to only those that are predicted to be functional in tissues relevant for the trait under consideration. On average across traits, the functional component for the top tissue as defined by FUN-LDA shows the highest enrichment relative to other methods, with approximately 2% of the SNPs (functional in the top tissue) explaining an estimated 30% of SNP heritability. Beyond the applications shown here, such functional predictions have numerous other applications. They can naturally be used in gene discovery studies to potentially improve power in sequence-based association tests such as SKAT and burden [67], and in fine-mapping studies [68, 69]. They can also be used in identifying regulatory regions that are depleted in functional variation in a specific tissue, similar to recent efforts to identify coding regions that are depleted in functional (e.g. missense, nonsense, and splice acceptor/donor variants) variation [7]. Other applications include improving power of trans-eQTL studies, by using the cell type and tissue specific functional predictions as prior information. Similarly, gene-gene and gene-environment interaction studies can benefit from an analysis focused on variants predicted to be functional in a cell type or tissue relevant to the trait under study.

Choosing the number of functional classes in the LDA model is not an easy task, partly because the number of functional classes is not well defined. We have focused here on a model with nine functional classes based on combining an objective measure such as the perplexity of the model and biological knowledge. There is some subjectivity in any method that seeks to partition the genome into functional classes, both in terms of the number of such classes and their interpretation. Further experiments that produce catalogs of specific types of elements with validated tissue-specific functions would aid in determining the number of states that a genomic annotation model should have, and the interpretation of those states, leading to potential improvements in the accuracy of such functional predictors. Such tissue-specific experimental data would also allow the use of supervised methods which could lead to improved tissue-specific functional scores.

Unlike our method, most of the existing segmentation methods smooth the genomic signal spatially. While they thereby use information from neighboring regions in making predictions for a particular variant, they may be less able to predict functionality of narrow regions with different histone modification profiles from neighboring regions. Another difference between our method and methods that use peak calls is that our method may be better able to integrate weak signals present in several histone marks for prediction. Furthermore, the use of the valley score allows our method to predict functional regions that are narrower in size compared to existing methods.

The Roadmap and ENCODE epigenomes mostly represent average epigenomes over distinct cell populations within a tissue, and it is unknown how the individual cell-types contribute to the average epigenomes. Such a bulk characterization undoubtedly conceals the complexity of epigenetic regulation. Investigation of epigenetic regulation at the single-cell level would provide a more detailed and accurate characterization of the function of variants in each cell. Although single-cell epigenomics data are currently scarce, with rapid technological development of single-cell methods such data should become available over the next few years, and the proposed methods can be readily applied in such settings.

We have computed FUN-LDA posterior probabilities for every position in the human genome for 127 tissue and cell types available in Roadmap. These scores are available at our website and can be imported into the UCSC Genome Browser. Note also that it is easy to make predictions in a new tissue once the model has been fit to the tissues in Roadmap. Furthermore, as with some other existing methods [14], it is possible to make predictions in a new tissue even if not all the epigenetic features we included are available, assuming one can impute the missing features by taking advantage of the correlations of epigenetic signals across both marks and samples as in ChromImpute [15].

## Acknowledgments

We gratefully acknowledge support by National Institutes of Health grants MH106910 (DB, ZH, II-L), AR065963 (LP, AC), DK105124 and the Herbert Irving Scholars Award (KK), MH100233 and the Seaver Foundation (JDB), the ATIP-Avenir program (VB). We thank Bin Xu and Badri Vardarajan for helpful discussions. We thank Andrew Brown for making the data on lead eQTLs in the Geuvadis and TwinsUK cohort available to us.

## WEB-BASED RESOURCES

1000 Genomes: http://www.1000genomes.org/

ChromHMM: http://compbio.mit.edu/ChromHMM/

ENCODE: https://www.encodeproject.org/

FUN-LDA: http://www.columbia.edu/~ii2135/funlda.html

GenoSkyline-Plus: http://genoSkyline.med.yale.edu/GenoSkyline

GTEx: http://www.gtexportal.org/home/

IDEAS: http://bx.psu.edu/~yuzhang/Roadmap_ideas/

Reg2Map:https://personal.broadinstitute.org/meuleman/reg2map/HoneyBadger2-intersect_release/

Roadmap Epigenomics: http://www.roadmapepigenomics.org/

Segway: http://noble.gs.washington.edu/proj/encyclopedia/

UCSC genome browser: https://genome.ucsc.edu/

### GWAS summary statistics

Age at menarche: http://www.reprogen.org/Menarche_Nature2014_GWASMetaResults_17122014.zip

Alopecia areata: http://www.broadinstitute.org/~sripke/share_links/sRSxpynHPaYRJ1SnYXD17eo3qK8IE6_daner_ALO4_1011b_mdsex/

Alzheimer’s disease: http://web.pasteur-lille.fr/en/recherche/u744/igap/igap_download.php

Autism: http://www.med.unc.edu/pgc/files/resultfiles/pgcasdeuro.gz

Bipolar Disorder: http://www.med.unc.edu/pgc/files/resultfiles/pgc.bip.2012-04.zip

BMI, Height: http://www.broadinstitute.org/collaboration/giant/index.php/GIANT_consortium_data_files

Coronary Artery Disease: ftp://ftp.sanger.ac.uk/pub/cardiogramplusc4d/cardiogram_gwas_results.zip

Crohn’s Disease: ftp://ftp.sanger.ac.uk/pub/consortia/ibdgenetics/cd-meta.txt.gz

Educational Attainment: http://ssgac.org/documents/SSGACRietveld2013.zip

Epilepsy: http://www.epigad.org/gwas_ilae2014/ILAE_All_Epi_11.8.14.txt.gz

Ever Smoked: http://www.med.unc.edu/pgc/files/resultfiles/tag.evrsmk.tbl.gz

Fasting Glucose: ftp://ftp.sanger.ac.uk/pub/magic/MAGIC_Manning_et_al_FastingGlucose_MainEffect.txt.gz

HDL: http://www.broadinstitute.org/mpg/pubs/lipids2010/HDL_ONE_Eur.tbl.sorted.gz

IGAN: dbGaP Study Accession: phs000431.v2.p1

LDL: http://www.broadinstitute.org/mpg/pubs/lipids2010/LDL_ONE_Eur.tbl.sorted.gz

Rheumatoid Arthritis: http://plaza.umin.ac.jp/yokada/datasource/files/GWASMetaResults/RA_GWASmeta_European_v2.txt.gz

Schizophrenia: http://www.med.unc.edu/pgc/files/resultfiles/scz2.snp.results.txt.gz

Triglycerides: http://www.broadinstitute.org/mpg/pubs/lipids2010/TG_ONE_Eur.tbl.sorted.gz

Type 2 Diabetes: http://www.diagram-consortium.org/downloads.html

Ulcerative Colitis: ftp://ftp.sanger.ac.uk/pub/consortia/ibdgenetics/ucmeta-sumstats.txt.gz

## 4. METHODS

### 4.1. LDA model for functional annotation

We propose an application of the latent Dirichlet allocation (LDA) model [24], a generative probabilistic model, in the setting of functional genomics annotations with the goal to compute posterior probabilities for variants to belong to different functional classes.

Let us assume that we have a set of m genetic variants in the training set, together with a set of k functional annotations. For each variant i, we have *k* tissue-specific functional scores: **Z_i_** = (*Z_i_*_1_,…, *Z_ik_*). Let **Z** = (**Z_1_**,…, **Z_m_**) be the set of (continuous) functional scores for all the variants. These scores are epigenetic features (histone modifications and DNase) from ENCODE and Roadmap Epigenomics across a varied set of tissues and cell types. Let *l* be the number of tissues, and *m_j_* be the number of variants with tissue j annotations in the training set (m = 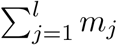) For each variant *i* ≤ *m* in the training set we denote by *t_i_* the corresponding tissue (i.e. the annotations corresponding to this variant are for tissue t_i_). For each tissue, the variants’ scores are represented as a mixture over latent functional classes, where each functional class is characterized by a distribution over variant scores. In what follows, for ease of presentation, we assume only two latent functional classes, but the number of classes can be chosen to be greater than two (see next section for a discussion on the choice of the number of functional classes). We let **C** = *(C_1_,…, C_m_)* denote the set of indicator variables for all the variants, where *C_i_* = 1 if variant i belongs to the first functional class and *C_i_* = 0 otherwise. We are not able to observe **C**.

Let ***α*** = (*α*_0_, *α*_1_) be the hyperparameter vector with *α*_0_, *α*_1_ > 0. We assume the functional annotation data has been generated from the following generative model:

1. For each tissue *j*, choose (1 – *π_j_*, *π_j_*) ~ Dir(*α*_0_, α_1_).
2. Given *π_j_*, for each variant *i* with t*_i_* = j choose a class *C_i_* ~ Bern(*π_j_*).
3. Given *C*_1_,*…, C*_m_, *Z_1_,…, Z_m_* are independently generated with each Z_*i*_ being generated from *F_1_* if *C_i_* = 1, and from F_0_ otherwise.

Here ***π*** = (*π*_1_,…, *π_l_*) and C are latent variables. We want to calculate the posterior probability for each variant i to be in the first functional class:

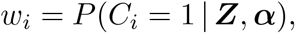

and the densities *f_0_* and *f_1_*. Also, we want to estimate the hyperparameter ***α*** = (*α*_0_, *α*_1_) empirically using Z. For a given tissue the conditional density of (*π*, *C*) given ***Z*** and ***α*** is:

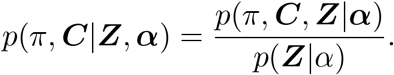

For the numerator we have:

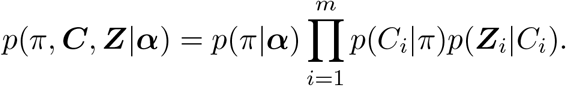

This is easy to compute. However the denominator is not. For the denominator we have:

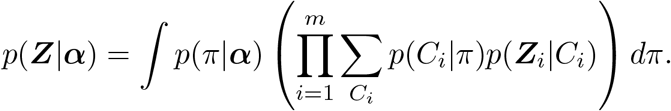

There are 2^*m*^ terms in the summation so this is difficult to compute for moderately large *m*. We propose instead to use a variational approach as described in [24]. In the variational inference approach we first introduce a family of distributions {*q*(·,·|***a***, ***w***)} over the latent variables (*π*, ***C***) with its own variational parameters ***a*** = (*a*_0_; *a*_1_) and *w* (these are tissue specific parameters).

Then

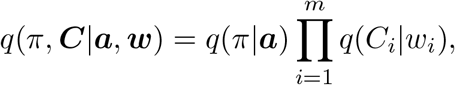

where *q*(*π*|***a***) is the density of Dir(***a***) and *q*(*C_i_*|*w_i_*) is the probability mass function of Bern(w_i_) for *i* = 1…m.

Using Jensen’s inequality we have:

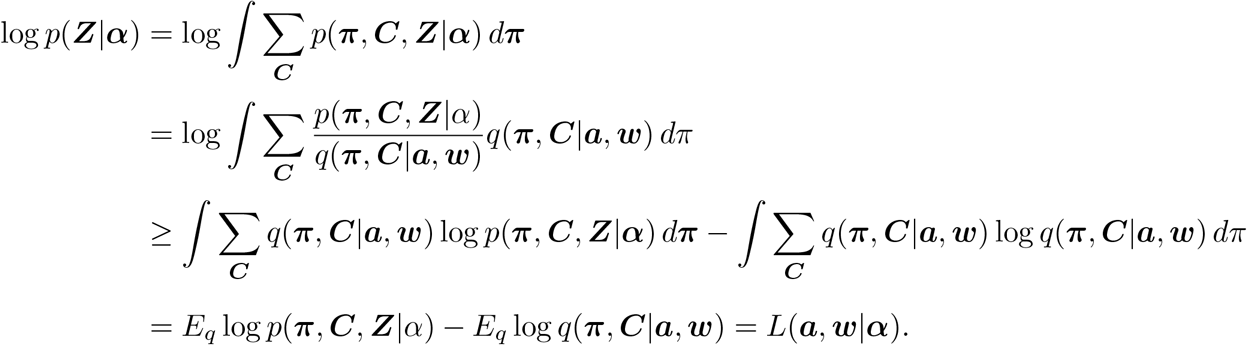

Note that *L*(***a*, *w***|***α***) is a lower bound on the log likelihood. So instead of maximizing the log likelihood directly we maximize this lower bound with respect to the variational parameters o and w, as well as the hyperparameter α. It can be shown that log *p*(***Z***|***α***) − *L*(***a, w***|***α***) is the Kullback-Leibler (KL) divergence between the true posterior *p*(***π, C***|***α, Z***) and the variational posterior *q*(***π, C***|***α, w***) with respect to *q*(***π, C***|***α, w***). Therefore by maximizing *L*(***a, w***|***α***) with respect to ***a*** and ***w***, we minimize the KL divergence between the variational posterior probability and the true posterior probability. Then we can estimate *P*(*C_i_* = 1|***α, Z***) by w_i_ for each variant i. Below we describe the variational inference algorithm.

**Variational Inference Algorithm.** Assume the initial state (*w*_1_,…, *w_m_*, *f*_0_, *f*_1_, ***α***). The algorithm proceeds as follows:

Step 1. *(Kernel Density Estimation)*

Fit a multivariate kernel density estimate for each annotation and component separately: 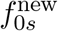 and 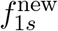 for each annotation *s* = 1,…,k, weighting variants by component membership probability. Specifically, for any **x** = (x_i_,…,x_k_) ∈ ℝ^*k*^ and s = 1,…,k, we let

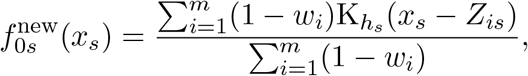

and

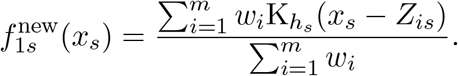

The scaled kernel K_*h*_*s*__ = 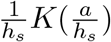, where *K*(·) is taken to be the probability density function of a standard normal, and the bandwidth parameter *h_s_* is chosen to be

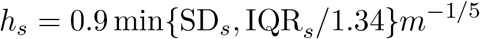

according to a rule of thumb due to Silverman [70], where SD_s_ and IQR_s_ are the standard deviation and interquartile range of annotation s, respectively. Then

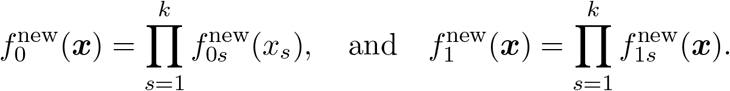

Step 2. *(Variational Step)*

For each tissue *j*, we obtain *w*_*i*_ for all variants *i* with *t_i_* = *j* and 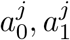 by maximizing the lower bound on the marginal likelihood of ***Z***, i.e. *L*(***a,w***|***α***), with respect to ***a*** and ***w***. Details are shown in the Supplemental Material.

This results in the following iterative algorithm:

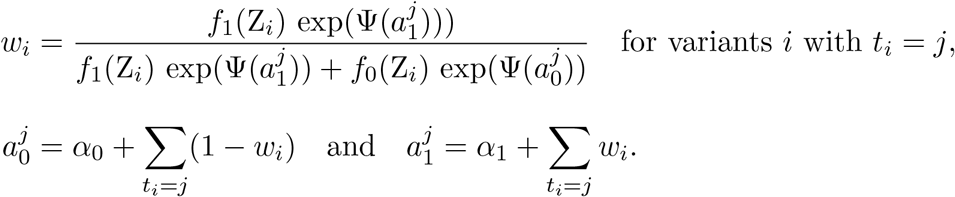

where Ψ(*x*) = *d* log Γ(*x*)/*dx* and Γ(*x*) is the Gamma function.

Step 3. *(Newton-Raphson algorithm to estimate the hyperparameters **α**)*

Obtain the empirical Bayes estimate of ***α*** = (*α*_0_; *α*_1_) by maximizing the bound *L*(***a***, ***w***|***α***) by using Newton-Raphson algorithm where ***a*** and ***w*** are from Step 2. That is, we find optimal ***α*** by iterating:

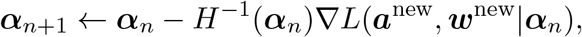

where *H*(***α***) is the Hessian matrix evaluated at current ***α***.

The gradient ∇*L*(***α***) has this form:

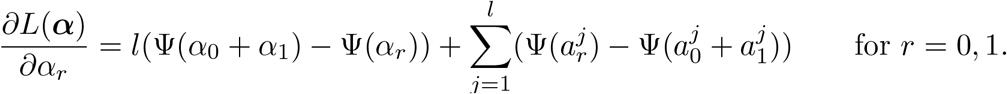

The Hessian matrix takes the following form:

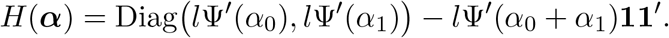

### 4.2. LDA implementation

We have implemented the above algorithm into an R package, FUN-LDA. In our implementation, we assume a symmetric Dirichlet prior, with *α* = 1, corresponding to a uniform distribution. For training purposes, we select 4000 random variants in each of the 127 tissues. The number of outer iterations in the variational inference algorithm is 250 and the number of inner iterations is 200.

FUN-LDA is computed by fitting the LDA model with nine classes to valley scores for the four activating histone modifications (H3K4me1, H3K4me3, H3K9ac, H3K27ac), and original DNase. For the histone modifications and DNase we start with negative log10 of the Poisson P-value of ChIP-seq or DNase counts relative to expected background counts, as output by ChromImpute [15]. The valley scores are computed as in [25]: for every window of 25 bp, we calculate the maximum score for the two regions from −100 to −500 bp and from 100 to 500 bp. If the score at the window of 25 bp is less than 90% of the minimum of those two maxima, we set the value in that window to that minimum. Otherwise, we set the value in that 25 bp window to 0. For each variant, we get a set of nine posterior probabilities for the variant to be in a specific functional class. To get a functional score, we sum the posterior probabilities for the active functional classes, namely ‘active promoters’ and ‘active enhancers’ (Supplemental Figure S2 and Table S3).

### 4.3. Prediction in a new tissue

Once the LDA model has been fit to the epigenetic data for cell types and tissues available in Roadmap, making predictions for a new cell type or tissue is easy. Basically, one only needs to run the iterative algorithm in Step 2 of the variational inference algorithm on the epigenetic data for the new tissue.

### 4.4. Choice of number of functional classes in the LDA model based on the perplexity measure

Choosing the number of functional classes in the LDA model is not straightforward. Too few classes can be insufficient and can lower the accuracy of the resulting classifier. Too many classes can lead to an overly complex model and is subject to overfitting.

Heuristic methods exist based on computing the perplexity of a model with a given number of clusters on held out datasets. Perplexity is used in information theory to describe how well a statistical model fits the data. The lower the perplexity, the better the model, and its generalization performance. In our case, if we let *L(**Z**_t_i__*) = log(*p*(***Z**_t_i__*| α)) be the log-likelihood for a held out set of variants for each tissue group *t_i_*, the perplexity is defined as

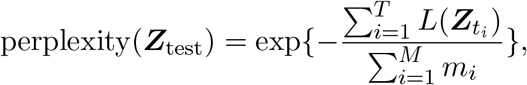

where T is the total number of tissues and m_i_ is the number of variants for tissue *t_i_*. Evaluating the perplexity measure directly is computationally intractable (the computation of the likelihood for each tissue involves a summation over *K^m_i_^* terms with K being the number of classes), and therefore we use the lower bound on the log-likelihood, i.e. *L*(***a***, ***w***|***α***) (see Supplemental Material), to derive an upper bound on the perplexity. This upper bound on the perplexity is referred to as the variational Bayesian bound on the perplexity. In the large data limit, the bound on the log perplexity evaluated on the training data converges to the Bayesian information criterion (BIC) for the model [71]. If the training and testing datasets are assumed to come from the same distributions, then the variational Bayesian bound on the log perplexity converges to the BIC.

### 4.5. Alternative functional annotation methods used in our comparisons

We compare our approach with the following state-of-the-art functional annotation methods.

#### 4.5.1. Individual histone modifications and DNase scores

Instead of integrating the various epigenetic marks, one can use the individual scores to predict functional variants. For the histone modifications and DNase we use negative log10 of the Poisson P-value of ChIP-seq or DNase counts relative to expected background counts, as output by ChromImpute [15]. In addition for DNase, we also use narrow peaks and gapped peaks (defined as broad peaks that contain at least one strong narrow peak).

#### 4.5.2. GenoSkyline - Multivariate Bernoulli mixture models

A simpler mixture model than the LDA described here is a two-component mixture model *ψ* = (*π*, *f*_0_, *f*_1_), where *f*_0_ and *f*_1_ are the probability densities for each of the components and n is a mixing parameter. We can fit such a model to data from each tissue separately, and calculate posterior probabilities for each variant to be in the ‘functional’ class given the observed scores Z, i.e. *Pψ*(*C_i_* = 1|**Z**). For tractability, it is often assumed that the individual scores are conditionally independent given the functional class. Such a two-component multivariate Bernoulli mixture model using dichotomized data from peak calling algorithms has been proposed in [23], an approach called GenoSkyline.

#### 4.5.3. ChromHMM

ChromHMM [14] is a method for chromatin state discovery and characterization by integrating multiple chromatin datasets. The underling algorithm is a multivariate Hidden Mixture Model that produces a segmentation of the genome; each segment is assigned a putative function based on enrichment analyses of different biological states in these segments. The ChromHMM 25-state model [15] is based on 12 marks, and, like ours, uses imputed data: H3K4me1, H3K4me2, H3K4me3, H3K9ac, H3K27ac, H4K20me1, H3K79me2, H3K36me3, H3K9me3, H3K27me3, H2A.Z and DNase. ChromHMM is based on complete pooling of data from multiple tissues and fitting a single model to this superdataset.

#### 4.5.4. Segway

Segway [16] is a genome segmentation approach, like ChromHMM, based on a dynamic Bayesian network (DBN) model. Segway is based on fitting separate models to data from each tissue. Segmentations for most of the cell types and tissues in Roadmap have been recently generated [72].

#### 4.5.5. IDEAS

IDEAS [20] is an integrative and discriminative epigenome annotation algorithm, that like ChromHMM and Segway, segments the genome and assigns each segment a specific functional class. Unlike ChromHMM and Segway, IDEAS models the correlations both along the genome and across cell types. Segmentations for all 127 cell types and tissues in Roadmap have been produced using IDEAS [21].

### 4.6. Generalized Jaccard index of overlap

We are interested in computing a similarity measure of predicted functional variants in two different tissues. Because the distribution of posterior probabilities in any one tissue is highly bimodal, with most of the mass at 0, and a small proportion of variants with posterior probabilities close to 1, in other words we are dealing with sparse binary data, a natural measure of similarity is the Jaccard measure of overlap, defined as follows. If **X** = (*x*_1_,…,*x*_*k*_) and **Y** = (*y*_1_,…,*y*_*k*_) are two vectors with *x_i_, y_i_* ≥ 0 (e.g. vectors of posterior probabilities for variants to be in the functional components for two different tissues), then the generalized Jaccard index of overlap is defined as:

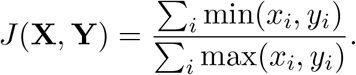

When **X** and **Y** are binary vectors, then the Jaccard index of overlap is simply the size of the intersection divided by the size of the union of the two sets. The closer it is to 1, the more overlap there is between the two sets. A Jaccard index of 0 means no overlap.

### 4.7. Promoter and tissue-specific enhancer regions

The promoter region of a protein-coding gene is defined as the union of the regions 2,500 bases upstream of any protein-coding transcripts for the gene, as defined by GENCODE version 24. For enhancer regions we use the Roadmap Stringent enhancer list available at the Reg2Map website.

### 4.8. eQTL enrichment

Let *G*_1_,…,*G*_44_ be the 44 GTEx tissues with at least 70 samples (Supplemental Table S4), and *R*_1;_…,*R_127_* be the 127 Roadmap tissues. For a given tissue in GTEx *G_i_* we are interested in identifying the Roadmap tissue *R_j_* with the highest enrichment in eQTLs from G_i_ relative to other tissues in Roadmap.

Let

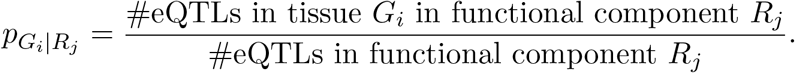

Note that the number of eQTLs in GTEx tissue *G_i_* is a weighted count, with an eQTL weighted by the inverse of the number of GTEx tissues in which the variant is eQTL, such that Σ_*i*_*pG_i_|R_j_* = 1.This way eQTLs that are unique to tissue Gi are given higher weight relative to eQTLs that are shared across many tissues. For GTEx tissue *G_i_*, to test whether there is an enrichment in the functional component of Roadmap tissue *R*_*j*_, we compare *p_Gi_*_|*Rj*_ with

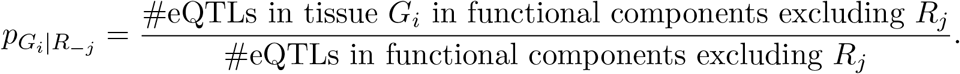

The null hypothesis is *H*_0_: *P*_*G_i_*|*R_j_*_ = *P*_*G_i_*|*R_−j_*_ vs. *H*_0_: *P*_*G_i_*|*R_j_*_ > *P*_*G_i_*|*R_−j_*_. We apply a two-sample proportion test for each Roadmap tissue *R_j_* and report the Roadmap tissue with minimum p value in Table 1.

The eQTLs that we used in these analyses are all significantly associated SNP-gene pairs in each of these 44 GTEx tissues, obtained using a permutation threshold-based approach as described by the GTEx Consortium [27]. For the follow-up study making use of eQTLs from Geuvadis and TwinsUK cohort, we use the lead eQTLs, i.e. those eQTLs most strongly associated with gene expression (publicly available for download from [28]).

### 4.9. Assessing pairwise correlations among 21 complex traits

Our aim here is to calculate a correlation matrix of 21 phenotypes based on the Z-scores from the LD score regression procedure, and a p value corresponding to each pair of phenotypes. From the LD score regression approach we obtain a matrix of Z-scores corresponding to 127 (p = 127) tissues and 21 (q = 21) phenotypes. The main issue we need to take into account when we compute the correlations and the p values is that the tissues are correlated.

Let *Z_ij_* be the *Z*-score corresponding to the i-th tissue and j-th phenotype; **Z_i_** = *(Z_i1_,…,Z_iq_*) and **Z^j^** = (*Z*_1*j*_,…,*Z_pj_*) be the row/column vectors of matrix **Z**. Since the elements of **Z** are Z-scores, we assume **Z_i_** ~ **N**(**0, Σ_q_**) and **Z^j^** ~ **N**(**0**, **Σ_p_**).

#### 4.9.1. Estimation of the correlation matrix

We aim to estimate **Σ_q_** but the problem is that **Z_i_**’s are not independent. To solve the problem, we propose the following perturbation method.

Let *B* be the number of perturbation replicates. For the b-th replicate, we generate *p* independent random variables from *N*(0, 1), *α_b_*_1_,…,*α_bp_*. Let

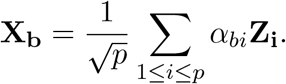

It can be shown that *cov*(**X_b_**) = **Σ_q_** and *cov*(**X_b_**, **X_b′_**) = 0 for any 1 ≤ *b*, *b*′ ≤ *B*. So we are able to use the uncorrelated perturbation samples **X_1_**,…, **X_B_** to approximate **Σ_q_** and the corresponding correlation matrix **P_q_**. We take *B* = 100, 000.

#### 4.9.2. P values corresponding to all pairs of phenotypes

For pairs from an uncorrelated bivariate normal distribution, the sampling distribution of a certain function of Pearson’s correlation coefficient follows Student’s t-distribution with degrees of freedom *M* — 2, where *M* is the number of uncorrelated random variables. Specifically, if the underlying variables have a bivariate normal distribution, the variable

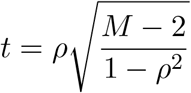

follows a Student’s t-distribution with degrees of freedom M — 2.

In our case, the number of uncorrelated random variables *M* depends on the correlation structure of the 127 tissues. M can be understood as the “effective number of tissues”. Similar to the calculation of “number of effective tests” by [73], we estimate M by applying an eigen-decomposition to the Jaccard matrix. Suppose λ_1_ ≥ λ_2_ ≥ ⋯ ≥ λ_*p*_ are the eigenvalues arranged in descending order. We estimate M by the smallest value such that 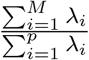>*C*. It should be noted that a smaller C will result in more conservative p values as the number of “effective tissues” is smaller, e.g. M = 124 when C = 99.5%, M = 96 when C = 95%. Too large or too small threshold C may cause M to be either overly liberal or overly conservative. The p values were calculated based on C = 99.5%.

### 4.10. Code availability

We have implemented the LDA algorithm into an R package, FUN-LDA. The package is available at the Comprehensive R Archive Network (CRAN): https://cran.r-project.org/web/packages/FUNLDA.

## Supplemental Material

### Stratified LD score regression approach to identify the tissue of interest

The stratified LD score regression approach [1] uses two sets of SNPs, reference SNPs and regression SNPs. The regression SNPs are SNPs that are used in a regression of *χ*^2^ statistics from GWAS studies against the “LD scores” of those regression SNPs. The LD score of a regression SNP is a numeric score which captures the amount of genetic variation tagged by the SNP. Here, following [1] we use as regression SNPs HapMap3 SNPs, chosen for their high imputation quality, and as reference SNPs those SNPs with minor allele count greater than 5 in the 379 European samples from the 1000 Genomes Project [2]. We first compute tissue-specific scores using each of our methods for the 9, 254, 335 SNPs with minor allele count greater than 5 in the 379 European samples from the 1000 Genomes Project, which we will subsequently use as our “reference SNPs” for LD score regression.

In the stratified LD score regression approach, a linear model is used to model a quantitative phenotype y_i_ for an individual *i*:

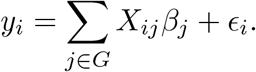

Here *G* is some set of SNPs, *X_ij_* is the standardized genotype of individual *i* at SNP *j*, *β_j_* is the effect size of SNP *j*, and ∈_*i*_ is mean-zero noise. In this framework, *β*, the vector of all the *β_j_*, is modeled as a mean-0 random vector with independent entries, and the variance of *β_j_* depends on the functional categories included in the model. We have a set of functional categories *C*_1_,*…,C_C_*, and the variance of a SNP’s effect size will depend on which functional categories it belongs to:

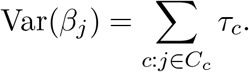

Here *τ*_*c*_ is the per-SNP contribution to heritability of SNPs in category *C_c_*. In [1], the authors show that under this model *τ*_c_ can be estimated through the following equation:

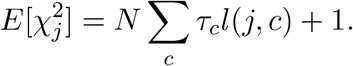

Here 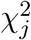 is the chi-squared statistic for SNP *j* from a GWAS study, *N* is the sample size from that study, and *l(j, c)* is the LD score of SNP j with respect to category 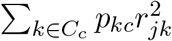. This equation therefore allows for the estimation of the T_*c*_ via the regression of the chi-squared statistics from a GWAS study on the LD scores of the regression SNPs.

Here, we extend the stratified LD score by allowing SNPs to be assigned to a category *C_c_* probabilistically, that is, we assume a probability *p_kc_* that SNP k belongs to category *C_c_*, and therefore that the variance of its effect size is affected by its membership in that category. This only involves minor changes to the above equations, namely, we have that

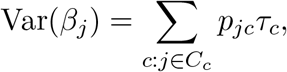

where *p_jc_* is the probability that SNP *j* belongs to category *C_c_*, and as above

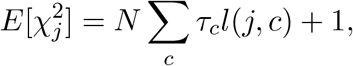

although now 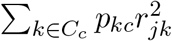 being the probability that SNP k belongs to category C_c_. We can therefore still estimate the t_c_ via the regression of the chi-squared statistics from a GWAS study on the LD scores of the regression SNPs, but in calculating these LD scores we weight the squared correlation of a SNP k with a regression SNP j by the probability that SNP k belongs to a particular category.

For each tissue and phenotype, and each of our functional scores, we fit a separate LD score regression model, including the LD score derived using the posterior probability that each regression SNP is in the functional component in that tissue, to estimate the per-SNP contribution of SNPs that belong to that component to heritability. To control for overlap of the tissue-specific functional score with other functional categories, we use the same 54 baseline categories used in [1], which represent various non-tissue-specific annotations, including histone modification measurements combined across tissues, measurements of open chromatin, and super enhancers.

### Summary of results for six SNPs in the literature, with evidence of regulatory function

– rs12350739 has been shown to influence human skin color by regulating transcription of nearby *BNC2* pigmentation gene [58]. In Supplemental Figure S6 we show the predictions for Roadmap tissue E059: Foreskin Melanocyte Primary Cells skin01, the tissue we deemed closest to the one used in the functional study, melanocyte cell lines.
– rs12740374: In [59] the authors show using human-derived hepatocytes that SNP rs12740374 creates a C/EBP (CCAAT/enhancer binding protein) transcription factor binding site and alters the hepatic expression of the SORT1 gene. In Supplemental Figure S7 we show the predictions for Roadmap tissue E066: Liver, the tissue we deemed closest to the one used in the functional study, human-derived hepatocytes.
– rs356168: In [60], the authors performed allele-specific TaqMan^®^ qRT-PCR analysis in human induced pluripotent stem cells (hIPSC)-derived neurons and show that this SNP regulates the expression of the *SNCA* gene, a gene implicated in the pathogenesis of Parkinson’s disease. In Supplemental Figure S8 we show the predictions for Roadmap tissue E007: H1 Derived Neuronal Progenitor Cultured Cells, the tissue we deemed closest to the one used in the functional study, hIPSC-derived neurons.
– rs2473307: In [61], the authors showed evidence that this SNP, associated with schizophrenia, reduces expression of *CDC42* gene in a human neuronal cell line. In Supplemental Figure S9 we show the predictions for Roadmap tissue E007, H1 Derived Neuronal Progenitor Cultured Cells.
– rs227727: In [62], the authors show that this SNP, in perfect LD with the most significant GWAS variant, alters the function of an enhancer. In Supplemental Figure S10, we show the predictions for Roadmap tissue E119, HMEC Mammary Epithelial Primary Cells.
– rs144361550: In [63], the authors show that this SNP, in strong LD with a lead GWAS variant, displays allele-specific transcriptional activity in primary melanocytes. Furthermore, mass spectrometry analyses using melanoma cell line revealed that RECQL is an unequivocal allele-preferential binder of rs144361550. In Supplemental Figure S11, we show the predictions for Roadmap tissue E059: Foreskin Melanocyte Primary Cells skin01, the tissue we deemed closest to the one used in the functional study, melanocyte cell lines.

### Inference and parameter estimation in the variational inference procedure

It can be shown that for a single tissue the lower bound on the log likelihood can be written as

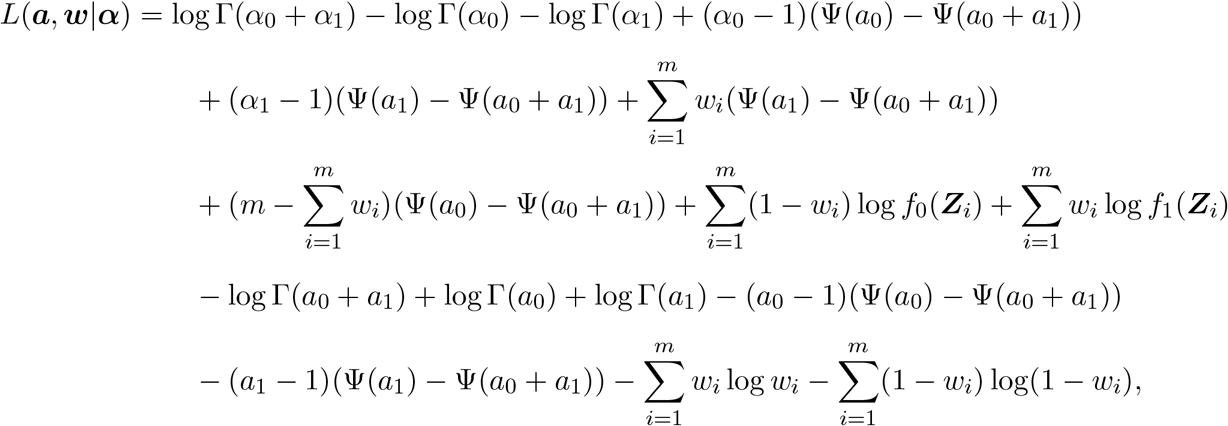

where Ψ(*x*) = *d* log Г(*x*)/*dx*

Maximizing *L*(***a***, ***w***|***α***) with respect to ***a*** and ***w***, respectively, we get

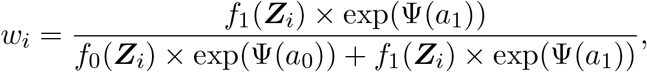

and

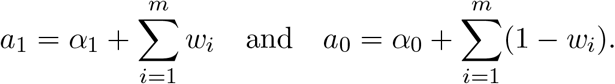

Given the optimal estimates of ***a*** and ***w***, we maximize the lower bound *L*(***a***, ***w***|***α***) with respect to the hyperparameter ***α*** by using the Newton-Raphson method as in [3]. Namely, we update ***α*** by iterating:

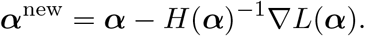

where the gradient ∇*L*(***α***) is:

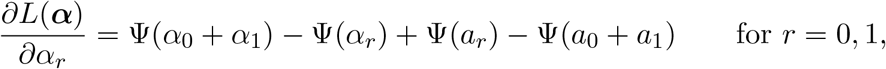

and for the Hessian matrix we have:

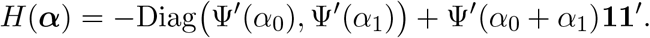

---

**Figure S1.**
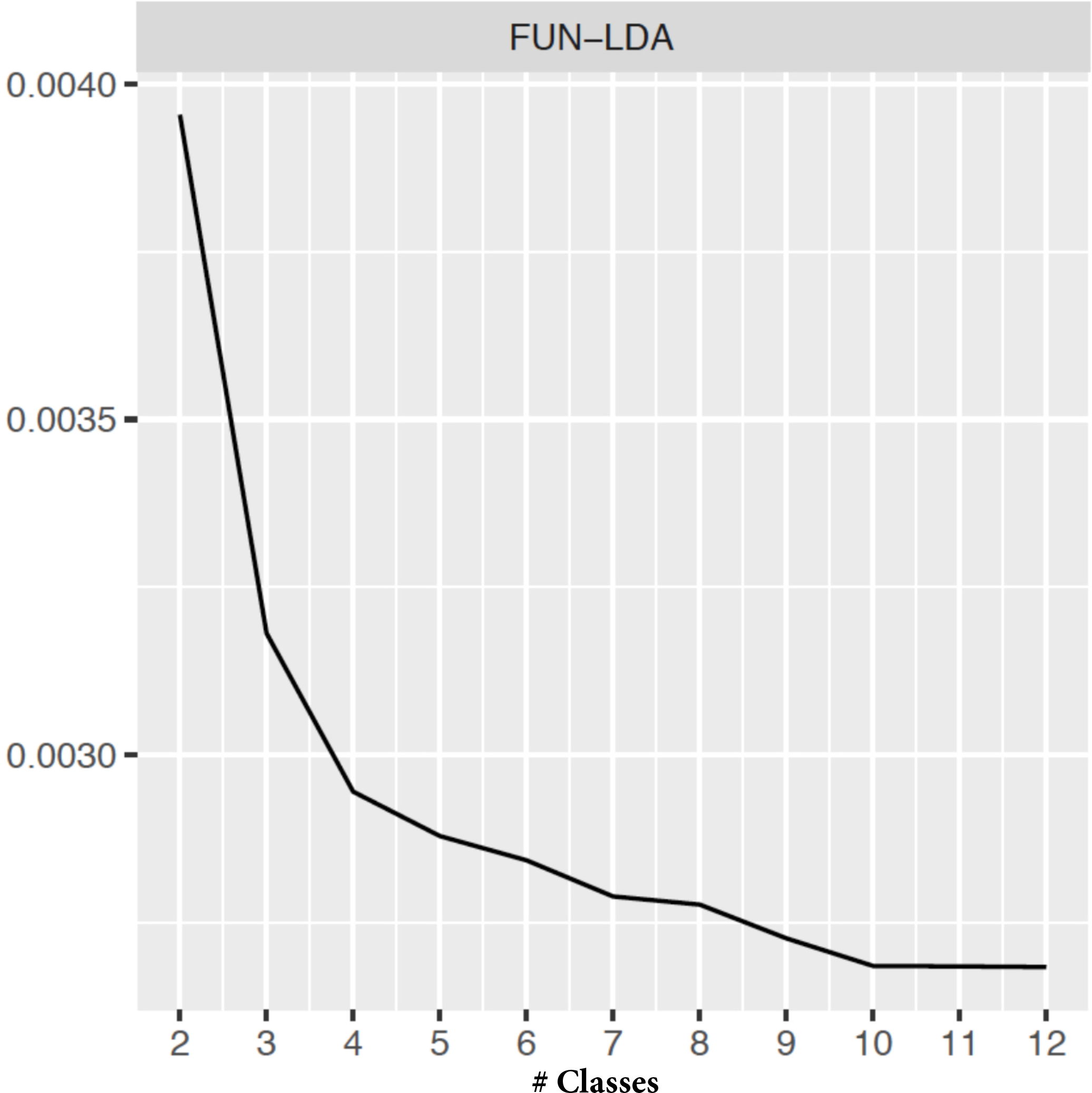
Perplexity measure of FUN-LDA models as a function of the number of classes.

**Figure S2.**
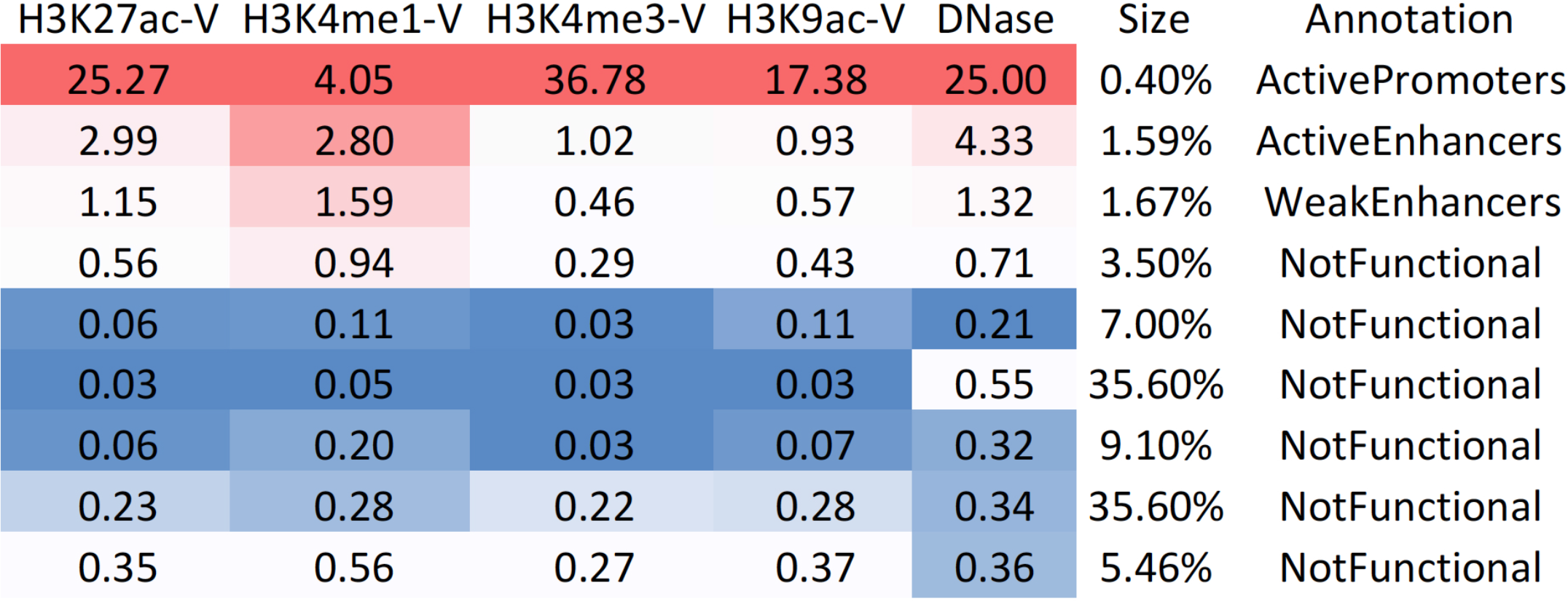
Heatmap of epigenetic features vs. class in the FUN-LDA model with nine classes across tissues and cell types in Roadmap.

**Figure S3.**
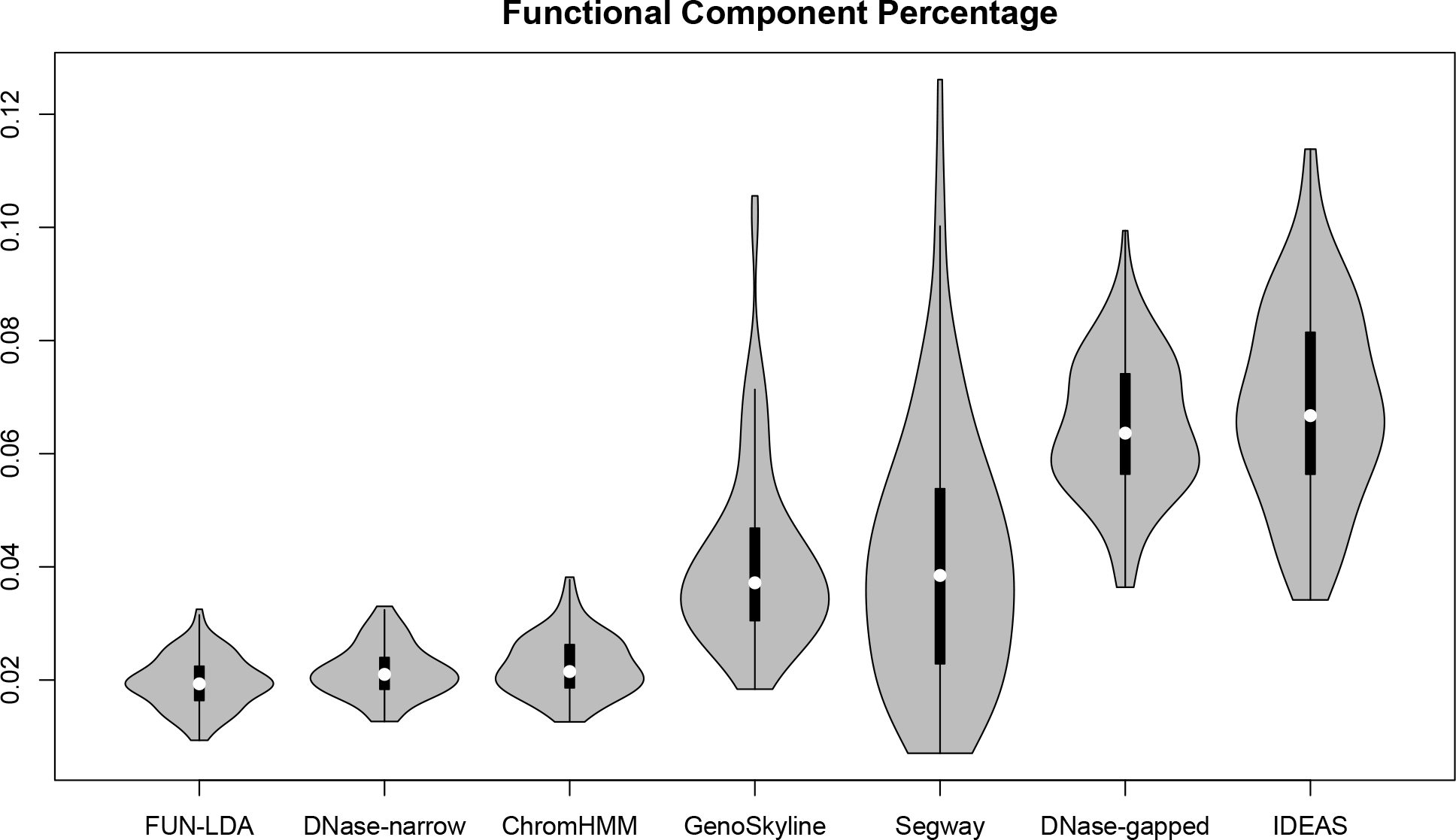
Violin plots showing the distribution of proportion of functional variants across tissues in Roadmap for each of several methods.

**Figure S4.**
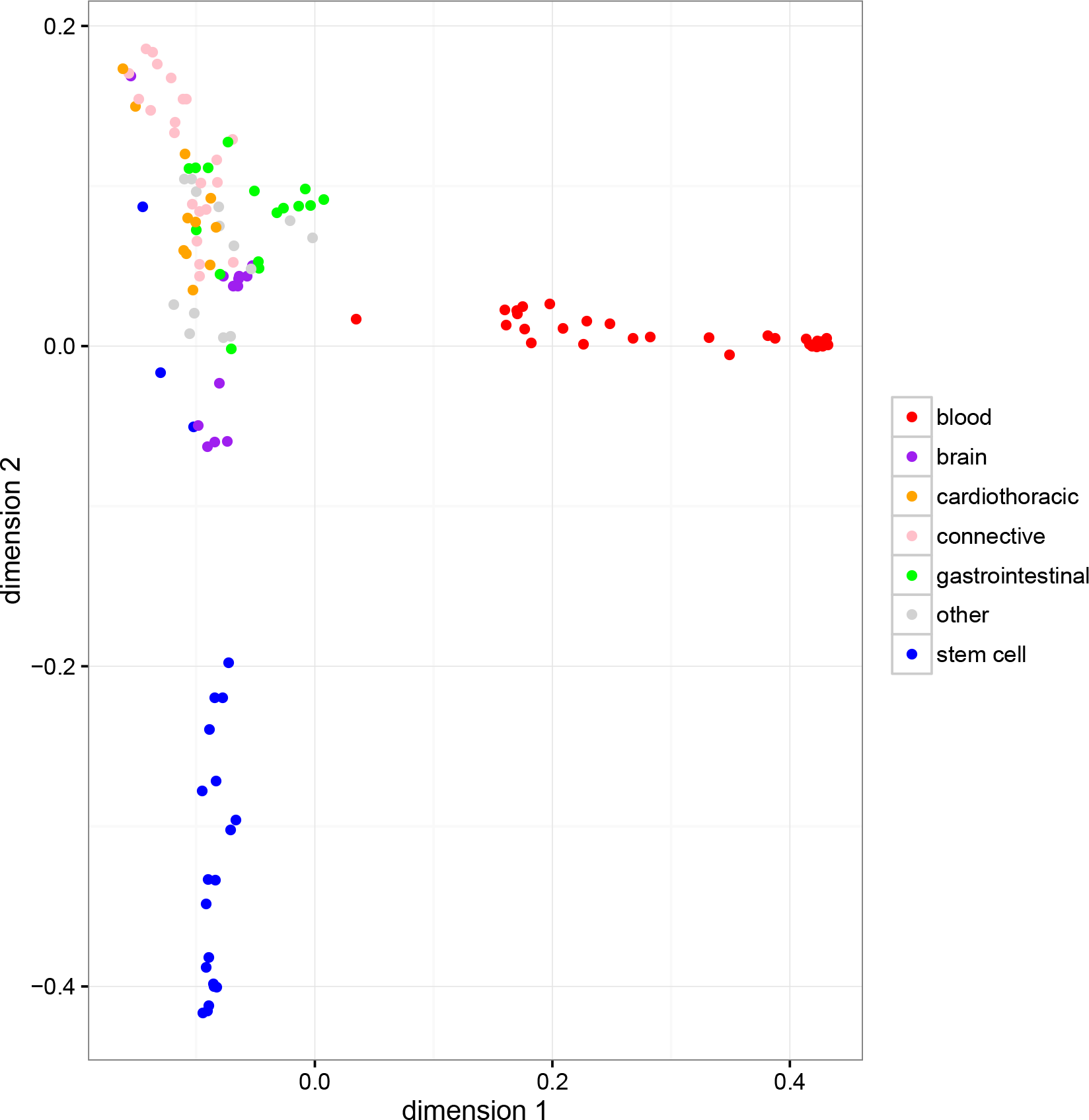
Multidimensional scaling plot of the correlations between the functional scores for the different tissues (FUN-LDA).

**Figure S5.**
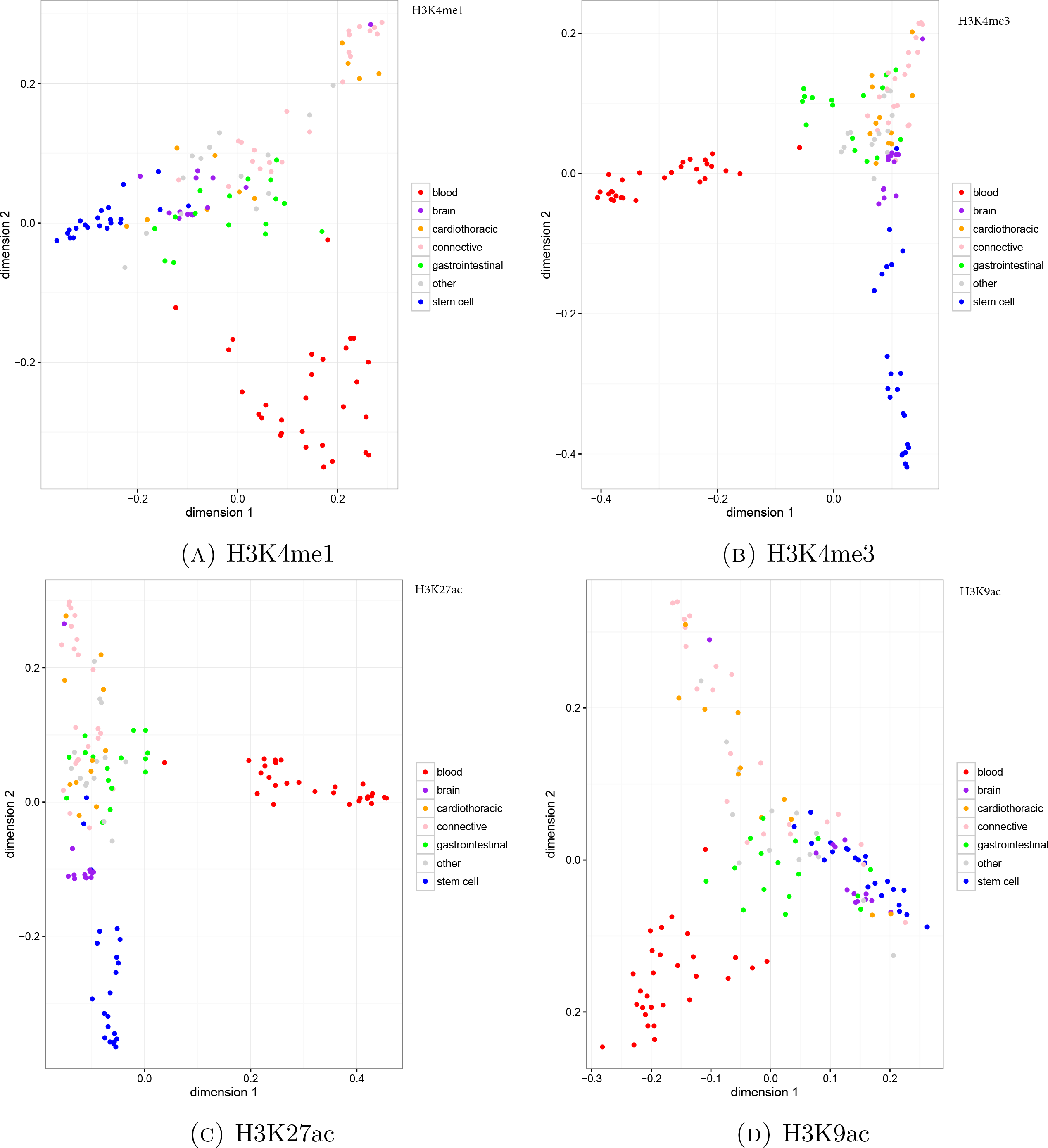
Multidimensional scaling plots of the correlations between the functional scores for the different tissues using individual histone marks.

**Figure S6.**
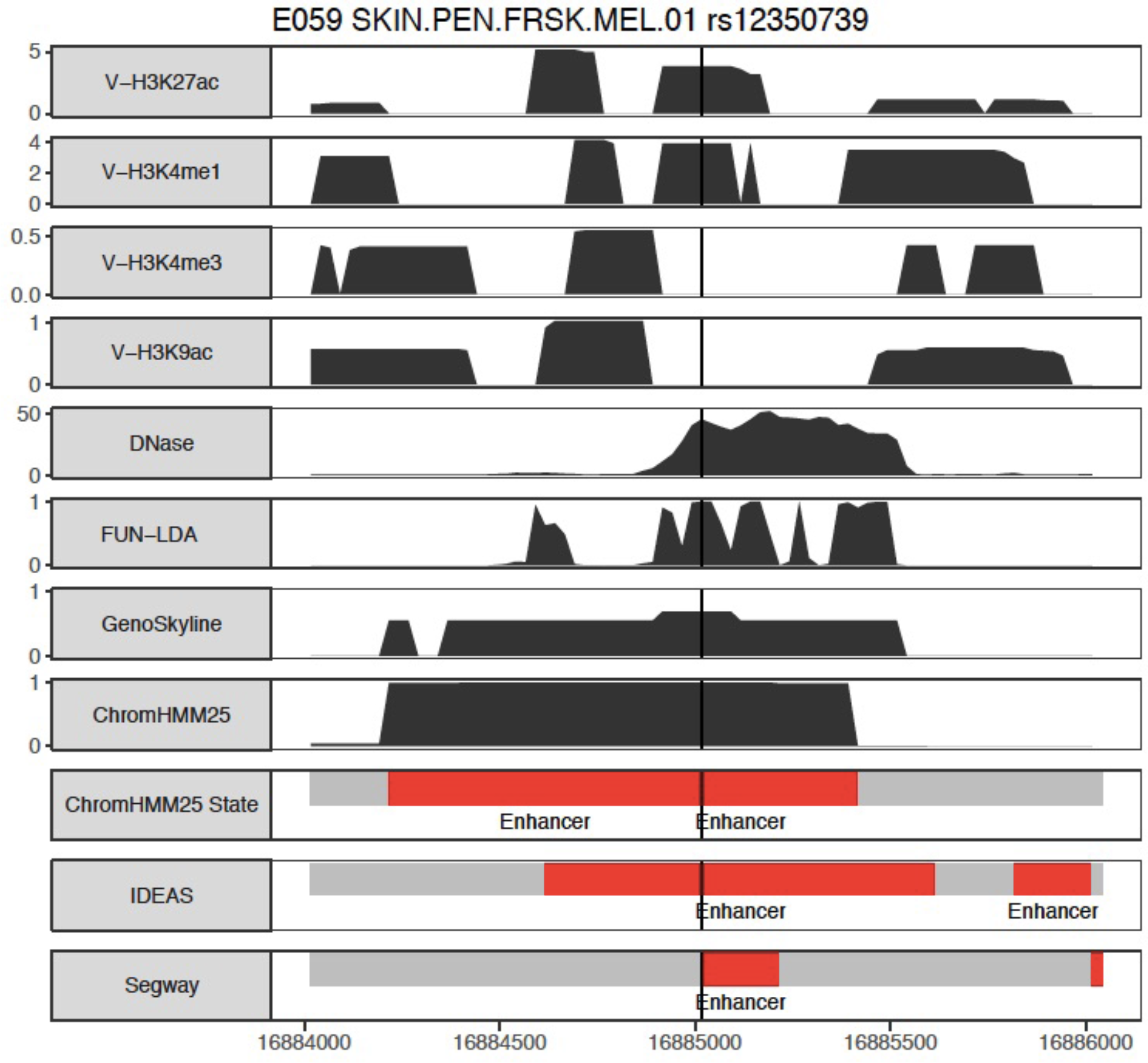
rs12350739 in Roadmap tissue E059. Valley scores for four activating histone marks and DNase, posterior probabilities from FUN-LDA, GenoSkyline, and ChromHMM (25 state model), and segmentations from ChromHMM, IDEAS and Segway are shown in 2 kb windows centered around the lead SNPs. For clarity we only highlight in the segmentations the type of states we consider functional (enhancer states in red, promoter states in blue) for the different segmentation approaches.

**Figure S7.**
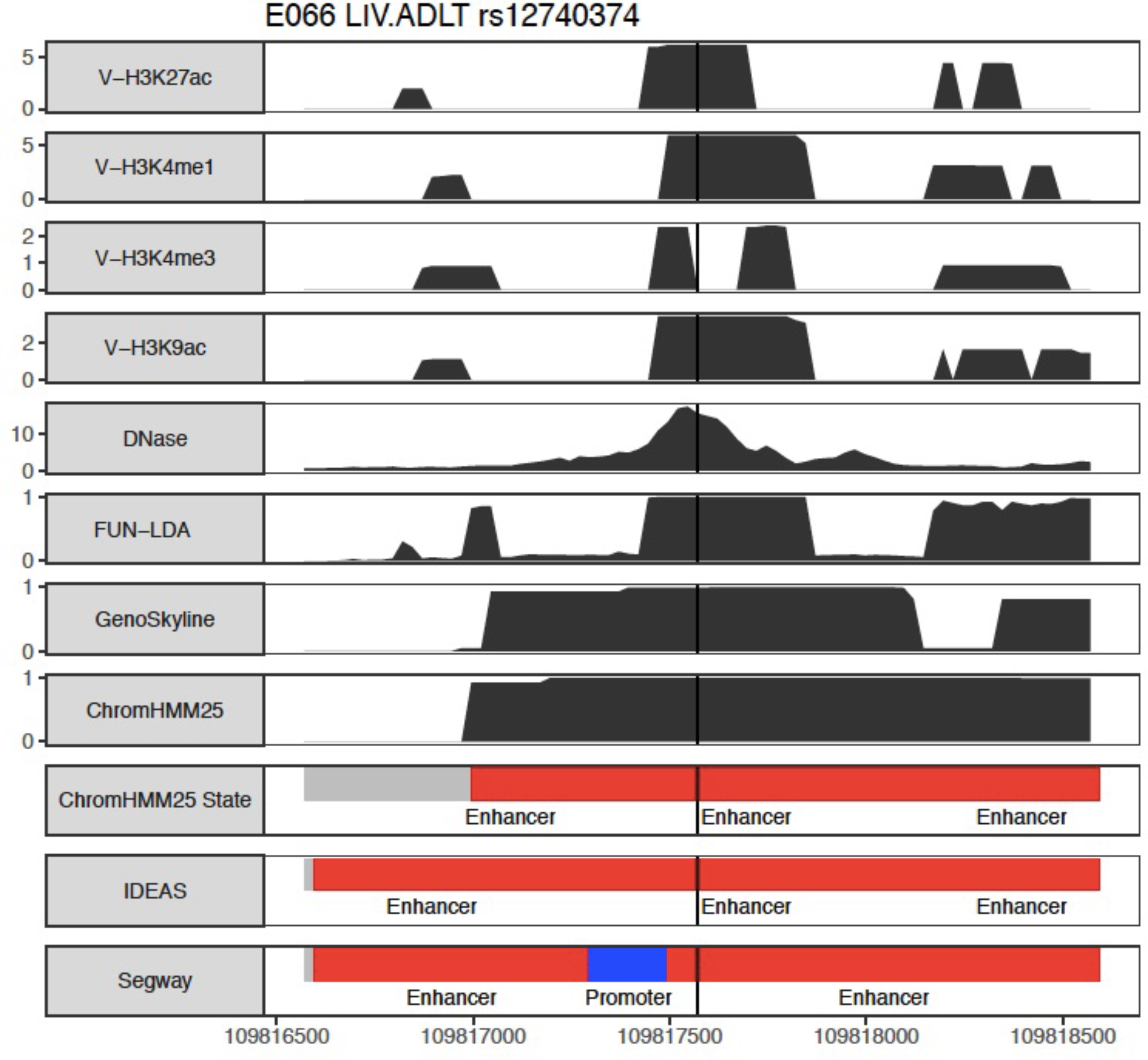
rs12740374 in Roadmap tissue E066. Valley scores for four activating histone marks and DNase, posterior probabilities from FUN-LDA, GenoSkyline, and ChromHMM (25 state model), and segmentations from ChromHMM, IDEAS and Segway are shown in 2 kb windows centered around the lead SNPs. For clarity we only highlight in the segmentations the type of states we consider functional (enhancer states in red, promoter states in blue) for the different segmentation approaches.

**Figure S8.**
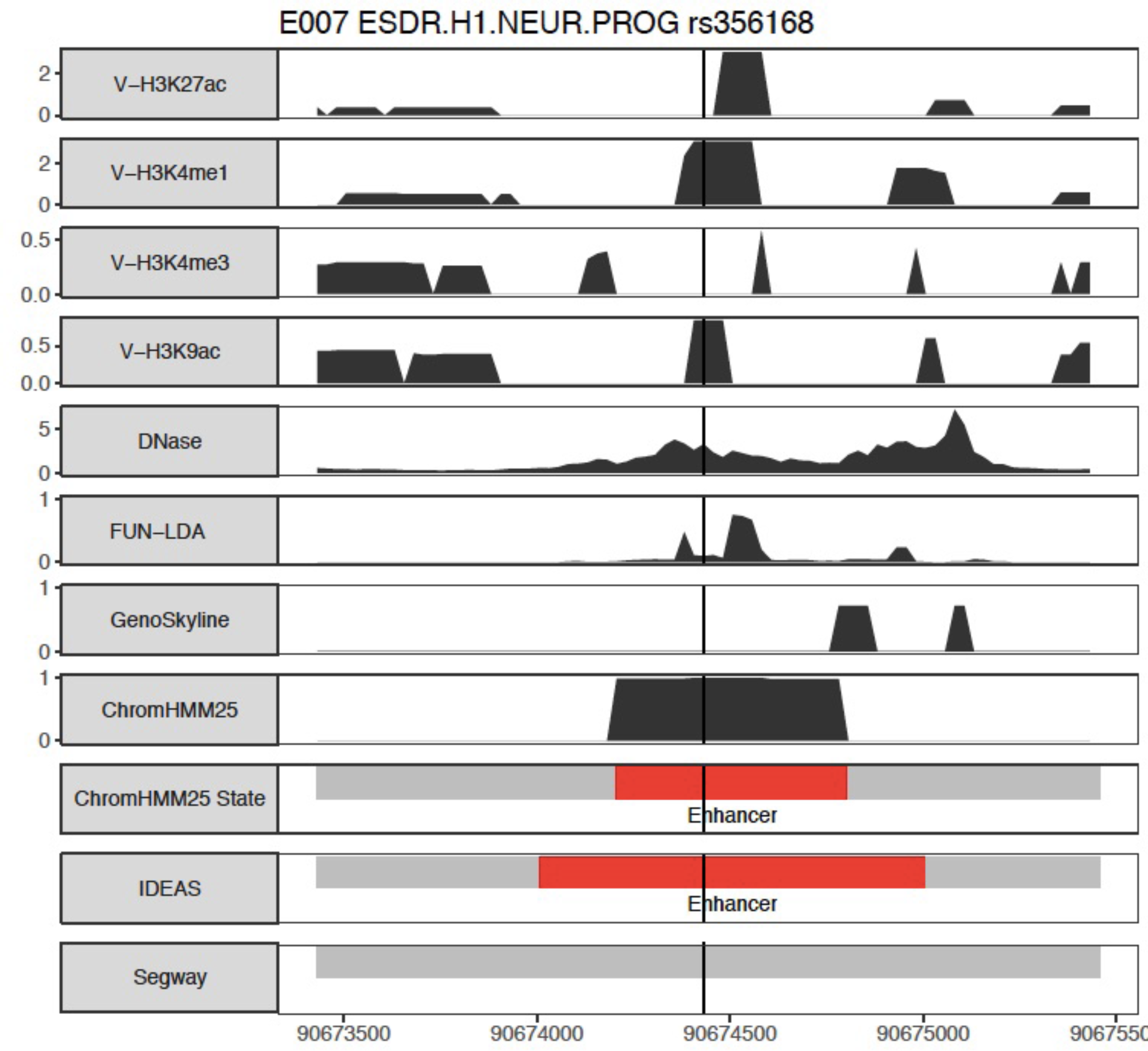
rs356168 in Roadmap tissue E007. Valley scores for four activating histone marks and DNase, posterior probabilities from FUN-LDA, GenoSkyline, and ChromHMM (25 state model), and segmentations from ChromHMM, IDEAS and Segway are shown in 2 kb windows centered around the lead SNPs. For clarity we only highlight in the segmentations the type of states we consider functional (enhancer states in red, promoter states in blue) for the different segmentation approaches.

**Figure S9.**
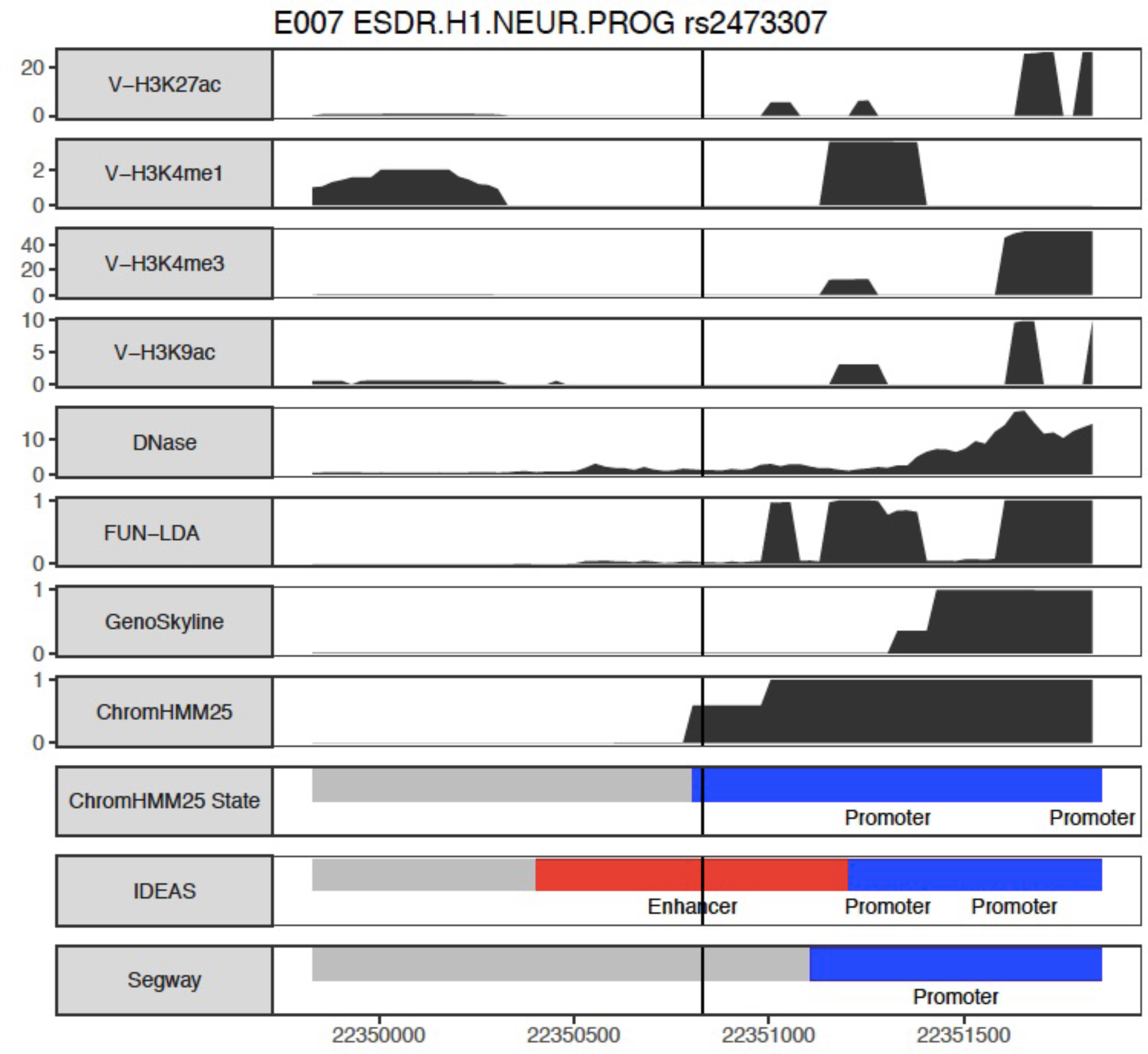
rs2473307 in Roadmap tissue E007. Valley scores for four activating histone marks and DNase, posterior probabilities from FUN-LDA, GenoSkyline, and ChromHMM (25 state model), and segmentations from ChromHMM, IDEAS and Segway are shown in 2 kb windows centered around the lead SNPs. For clarity we only highlight in the segmentations the type of states we consider functional (enhancer states in red, promoter states in blue) for the different segmentation approaches.

**Figure S10.**
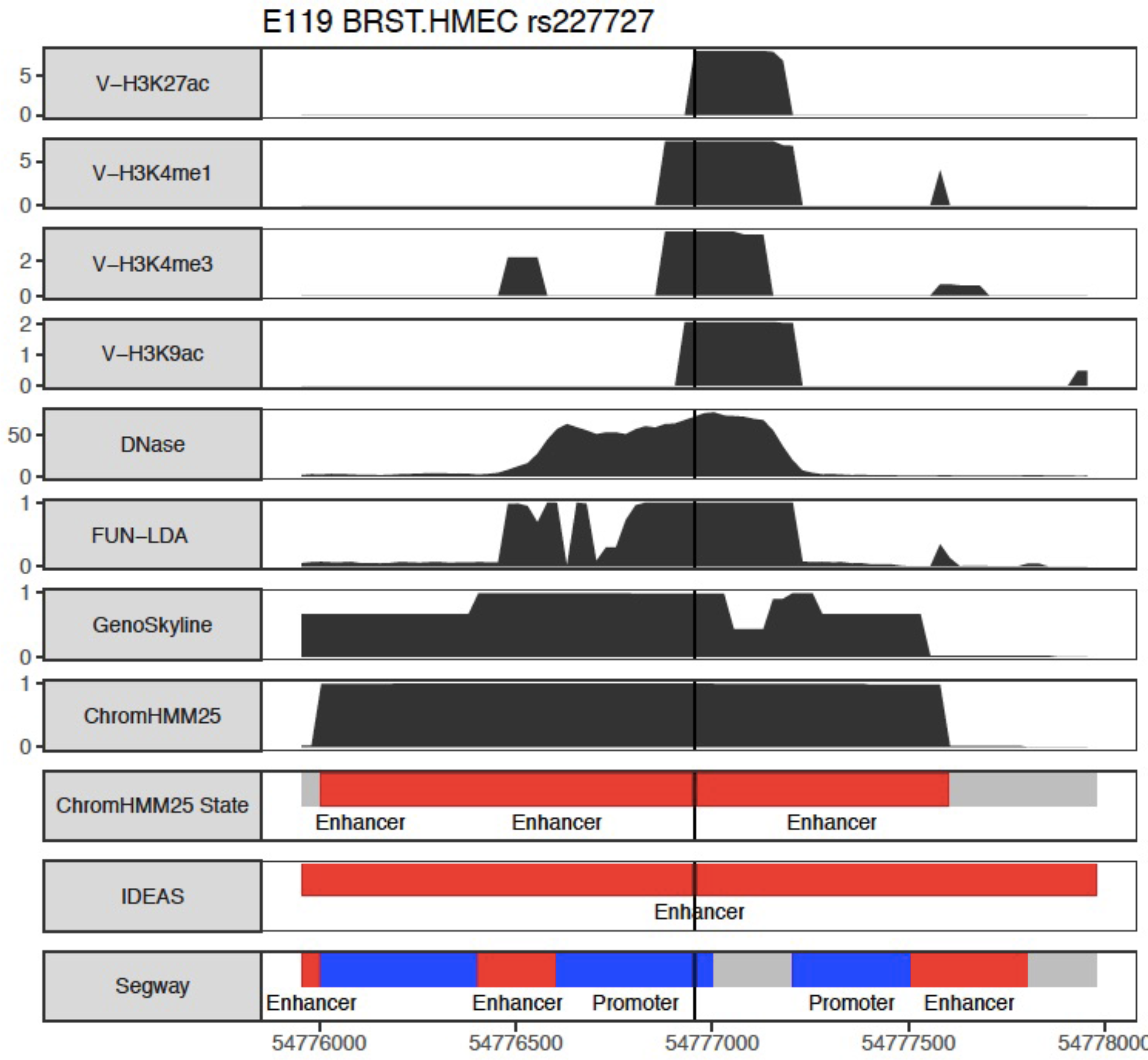
rs227727 in Roadmap tissue E119. Valley scores for four activating histone marks and DNase, posterior probabilities from FUN-LDA, GenoSkyline, and ChromHMM (25 state model), and segmentations from ChromHMM, IDEAS and Segway are shown in 2 kb windows centered around the lead SNPs. For clarity we only highlight in the segmentations the type of states we consider functional (enhancer states in red, promoter states in blue) for the different segmentation approaches.

**Figure S11.**
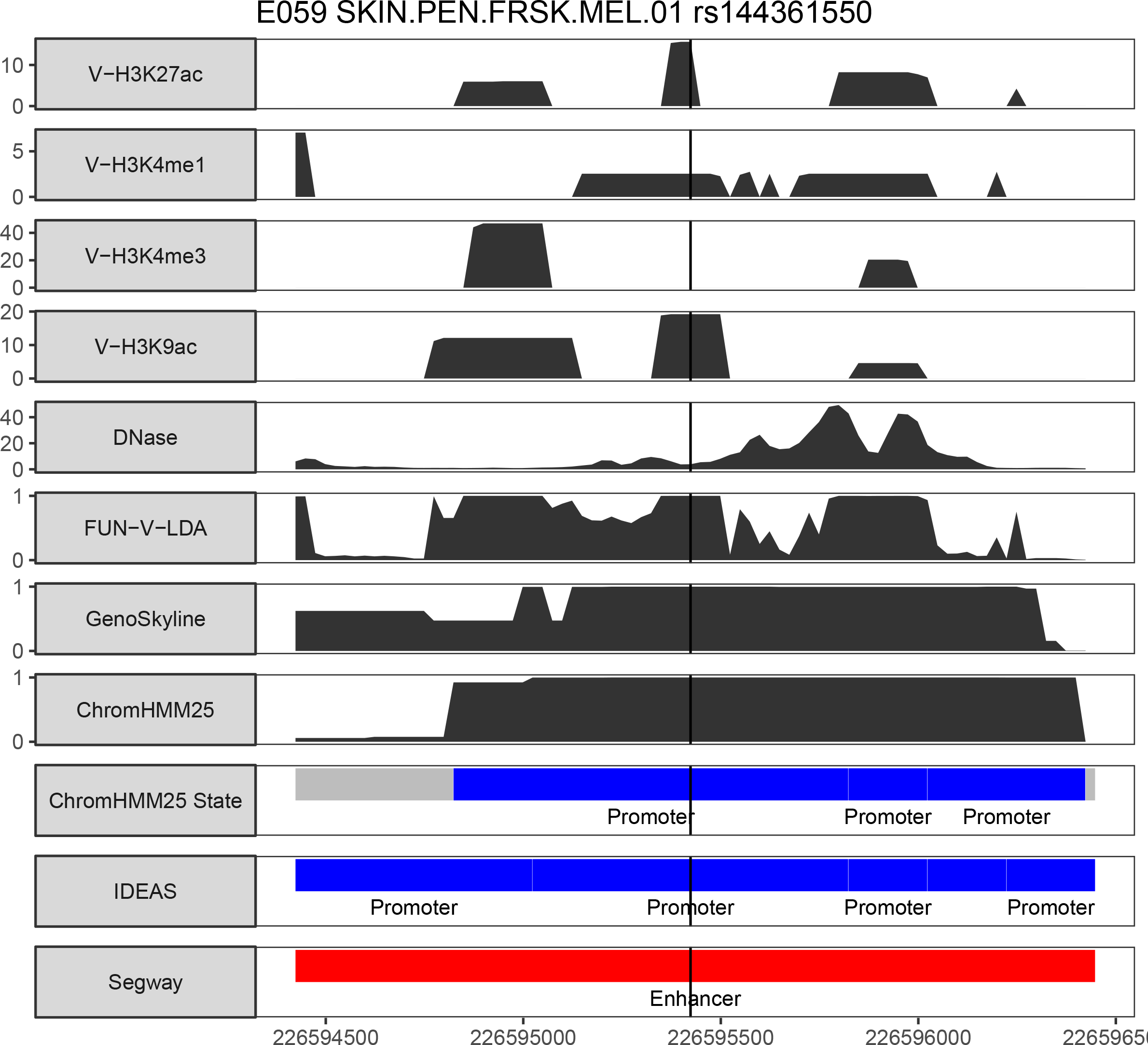
rs144361550 in Roadmap tissue E059. Valley scores for four activating histone marks and DNase, posterior probabilities from FUN-LDA, GenoSkyline, and ChromHMM (25 state model), and segmentations from ChromHMM, IDEAS and Segway are shown in 2 kb windows centered around the lead SNPs. For clarity we only highlight in the segmentations the type of states we consider functional (enhancer states in red, promoter states in blue) for the different segmentation approaches.

**Figure S12.**
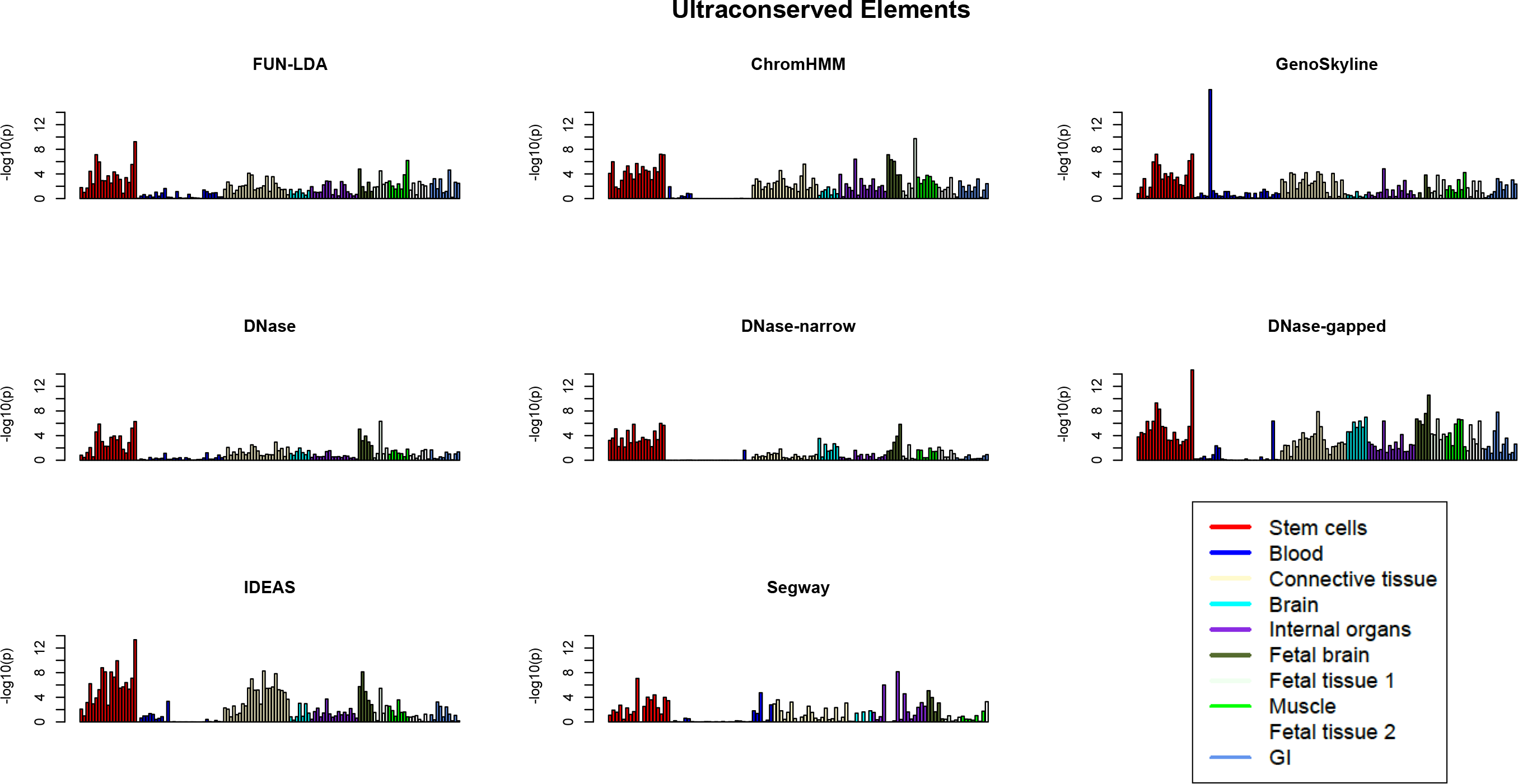
For each of several functional scores and across cell types and tissues in Roadmap, the p values from Wilcoxon rank sum test, comparing the ranks of functional scores for variants in validated enhancers in ultra conserved sequence elements vs. the ranks for the remaining variants in ultra conserved sequence elements are reported. The different tissues are grouped into several types (Supplemental Table S11).

**Table s1:**
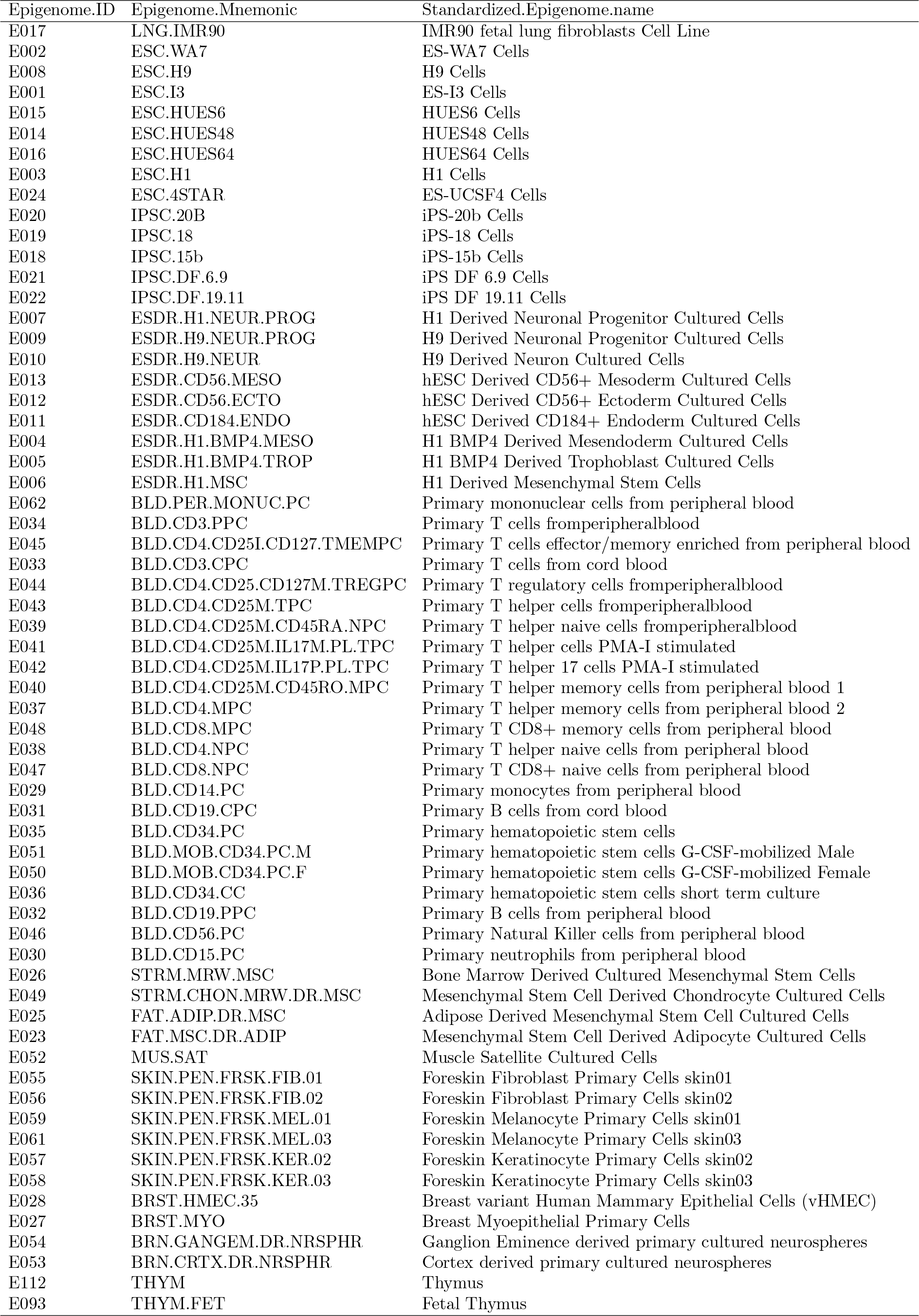
Tissues and Cell Types in Roadmap (part 1)

**Table s2:**
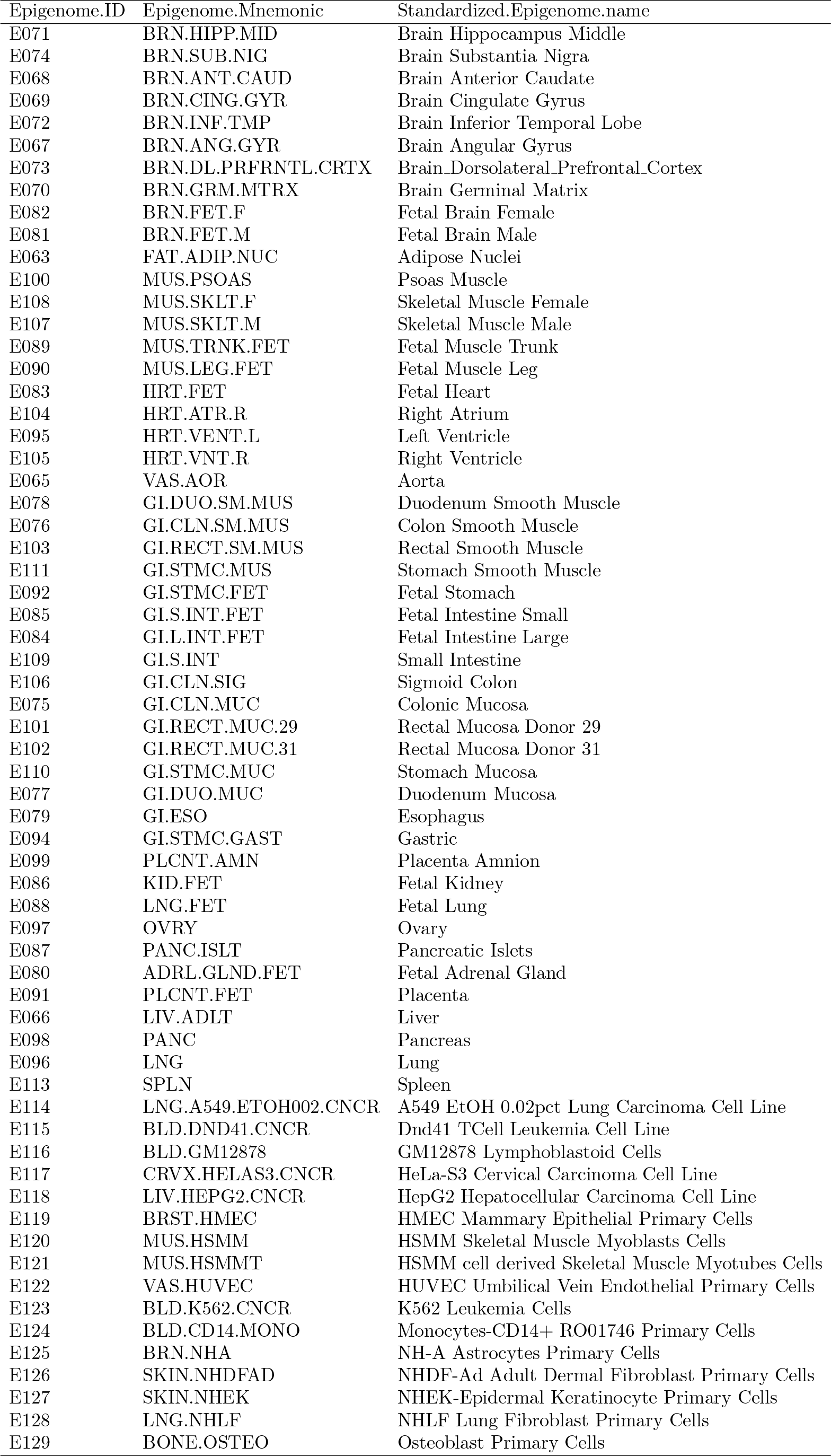
Tissues and Cell Types in Roadmap (part 2)

**Table s3:** Definition of the functional class for the five integrative methods considered.

| Method | Functional Class Definition |
| --- | --- |
| FUN-LDA | States 1 and 2 (active promoters and enhancers) in Supplemental Figure S2 |
| GenoSkyline | The functional class as defined in [23] |
| ChromHMM (25 state model) | 1_TssA, 2_PromU, 3_PromD1, 4_PromD2, 13_EnhA1, 14_EnhA2, 15_EnhAF |
| Segway | Promoters and Enhancers [72] |
| IDEAS | 4_Enh, 6_EnhG, 8_TssAFlnk, 10_TssA, 14_TssWk, 17_EnhGA [21] |

**Table s4:** GTEx tissues and sample sizes.

| Tissue | Sample size |
| --- | --- |
| Muscle - Skeletal | 361 |
| Whole Blood | 338 |
| Skin - Sun Exposed (Lower leg) | 302 |
| Adipose - Subcutaneous | 298 |
| Artery - Tibial | 285 |
| Lung | 278 |
| Thyroid | 278 |
| Cells - Transformed fibroblasts | 272 |
| Nerve - Tibial | 256 |
| Esophagus - Mucosa | 241 |
| Esophagus - Muscularis | 218 |
| Artery - Aorta | 197 |
| Skin - Not Sun Exposed (Suprapubic) | 196 |
| Heart - Left Ventricle | 190 |
| Adipose - Visceral (Omentum) | 185 |
| Breast - Mammary Tissue | 183 |
| Stomach | 170 |
| Colon - Transverse | 169 |
| Heart - Atrial Appendage | 159 |
| Testis | 157 |
| Pancreas | 149 |
| Esophagus - Gastroesophageal Junction | 127 |
| Adrenal Gland | 126 |
| Colon - Sigmoid | 124 |
| Artery - Coronary | 118 |
| Cells - EBV-transformed lymphocytes | 114 |
| Brain - Cerebellum | 103 |
| Brain - Caudate (basal ganglia) | 100 |
| Liver | 97 |
| Brain - Cortex | 96 |
| Brain - Nucleus accumbens (basal ganglia) | 93 |
| Brain - Frontal Cortex (BA9) | 92 |
| Brain - Cerebellar Hemisphere | 89 |
| Spleen | 89 |
| Pituitary | 87 |
| Prostate | 87 |
| Ovary | 85 |
| Brain - Putamen (basal ganglia) | 82 |
| Brain - Hippocampus | 81 |
| Brain - Hypothalamus | 81 |
| Vagina | 79 |
| Small Intestine - Terminal Ileum | 77 |
| Brain - Anterior cingulate cortex (BA24) | 72 |
| Uterus | 70 |
| Brain - Amygdala | 62 |
| Brain - Spinal cord (cervical c-1) | 59 |
| Brain - Substantia nigra | 56 |
| Minor Salivary Gland | 51 |
| Kidney - Cortex | 26 |
| Bladder | 11 |
| Cervix - Ectocervix | 6 |
| Fallopian Tube | 6 |
| Cervix - Endocervix | 5 |

**Table s5:** Results from stratified LD score regression for the different methods (part 1).

| Trait | Method | Roadmap Epigenome Name | -log10(p) |
| --- | --- | --- | --- |
| AgeAtMenarche | ChromHMM | Cortex derived primary cultured neurospheres | 4.31 |
| AgeAtMenarche | DNase | hESC Derived CD56+ Ectoderm Cultured Cells | 4.76 |
| AgeAtMenarche | DNase-gapped | iPS DF 6.9 Cells | 4.16 |
| AgeAtMenarche | DNase-narrow | ES-UCSF4 Cells | 7.36 |
| AgeAtMenarche | FUN-LDA | H9 Derived Neuron Cultured Cells | 6.15 |
| AgeAtMenarche | GenoSkyline | H1 Derived Neuronal Progenitor Cultured Cells | 7.96 |
| AgeAtMenarche | IDEAS | H1 Derived Neuronal Progenitor Cultured Cells | 3.47 |
| AgeAtMenarche | Segway | H1 Derived Neuronal Progenitor Cultured Cells | 9.91 |
| Alopecia | ChromHMM | Primary T helper cells PMA-I stimulated | 3.31 |
| Alopecia | DNase | Primary T helper 17 cells PMA-I stimulated | 2.10 |
| Alopecia | DNase-gapped | Primary T helper 17 cells PMA-I stimulated | 4.04 |
| Alopecia | DNase-narrow | Primary T helper memory cells from peripheral blood 1 | 3.81 |
| Alopecia | FUN-LDA | Primary T cells from cord blood | 3.90 |
| Alopecia | GenoSkyline | Primary T helper memory cells from peripheral blood 2 | 3.23 |
| Alopecia | IDEAS | Primary T helper 17 cells PMA-I stimulated | 4.48 |
| Alopecia | Segway | Primary T helper 17 cells PMA-I stimulated | 5.27 |
| Alzheimers | ChromHMM | Primary hematopoietic stem cells | 1.86 |
| Alzheimers | DNase | Monocytes-CD14+ RO01746 Primary Cells | 2.05 |
| Alzheimers | DNase-gapped | Primary hematopoietic stem cells G-CSF-mobilized Male | 3.96 |
| Alzheimers | DNase-narrow | Primary hematopoietic stem cells G-CSF-mobilized Male | 3.59 |
| Alzheimers | FUN-LDA | Primary hematopoietic stem cells G-CSF-mobilized Male | 3.78 |
| Alzheimers | GenoSkyline | Monocytes-CD14+ RO01746 Primary Cells | 2.91 |
| Alzheimers | IDEAS | Primary hematopoietic stem cells G-CSF-mobilized Male | 4.06 |
| Alzheimers | Segway | Primary hematopoietic stem cells G-CSF-mobilized Male | 3.79 |
| Autism | ChromHMM | Fetal Brain Female | 1.19 |
| Autism | DNase | Primary monocytes from peripheral blood | 1.64 |
| Autism | DNase-gapped | Primary monocytes from peripheral blood | 2.16 |
| Autism | DNase-narrow | Monocytes-CD14+ RO01746 Primary Cells | 1.94 |
| Autism | FUN-LDA | Primary monocytes from peripheral blood | 2.41 |
| Autism | GenoSkyline | Brain Dorsolateral Prefrontal Cortex | 1.26 |
| Autism | IDEAS | Liver | 2.54 |
| Autism | Segway | Monocytes-CD14+ RO01746 Primary Cells | 2.34 |
| BipolarDisorder | ChromHMM | Primary monocytes from peripheral blood | 2.27 |
| BipolarDisorder | DNase | Monocytes-CD14+ RO01746 Primary Cells | 2.23 |
| BipolarDisorder | DNase-gapped | Monocytes-CD14+ RO01746 Primary Cells | 3.48 |
| BipolarDisorder | DNase-narrow | Monocytes-CD14+ RO01746 Primary Cells | 2.48 |
| BipolarDisorder | FUN-LDA | Fetal Brain Female | 3.20 |
| BipolarDisorder | GenoSkyline | Psoas Muscle | 3.73 |
| BipolarDisorder | IDEAS | Fetal Brain Male | 3.30 |
| BipolarDisorder | Segway | Brain Dorsolateral Prefrontal Cortex | 3.70 |
| BMI | ChromHMM | Fetal Brain Female | 2.94 |
| BMI | DNase | ES-UCSF4 Cells | 1.12 |
| BMI | DNase-gapped | ES-UCSF4 Cells | 2.58 |
| BMI | DNase-narrow | ES-UCSF4 Cells | 4.29 |
| BMI | FUN-LDA | Brain Germinal Matrix | 4.79 |
| BMI | GenoSkyline | Brain Dorsolateral Prefrontal Cortex | 6.47 |
| BMI | IDEAS | Brain Angular Gyrus | 4.44 |
| BMI | Segway | iPS DF 19.11 Cells | 4.49 |
| CoronaryArteryDisease | ChromHMM | Liver | 3.38 |
| CoronaryArteryDisease | DNase | Liver | 2.62 |
| CoronaryArteryDisease | DNase-gapped | Liver | 4.67 |
| CoronaryArteryDisease | DNase-narrow | Lung | 3.51 |
| CoronaryArteryDisease | FUN-LDA | Liver | 4.61 |
| CoronaryArteryDisease | GenoSkyline | Lung | 4.25 |
| CoronaryArteryDisease | IDEAS | Adipose Nuclei | 3.65 |
| CoronaryArteryDisease | Segway | Small Intestine | 5.70 |

**Table s6:** Results from stratified LD score regression for the different methods (part 2).

| Trait | Method | Roadmap Epigenome Name | -log10(p) |
| --- | --- | --- | --- |
| CrohnsDisease | ChromHMM | Primary T helper 17 cells PMA-I stimulated | 6.39 |
| CrohnsDisease | DNase | Primary T helper cells PMA-I stimulated | 3.84 |
| CrohnsDisease | DNase-gapped | Primary B cells from peripheral blood | 6.89 |
| CrohnsDisease | DNase-narrow | Primary T helper 17 cells PMA-I stimulated | 6.90 |
| CrohnsDisease | FUN-LDA | Primary B cells from cord blood | 6.25 |
| CrohnsDisease | GenoSkyline | Primary Natural Killer cells from peripheral blood | 4.95 |
| CrohnsDisease | IDEAS | Primary T helper memory cells from peripheral blood 1 | 7.60 |
| CrohnsDisease | Segway | Primary T helper 17 cells PMA-I stimulated | 7.53 |
| EducationalAttainment | ChromHMM | Fetal Brain Female | 4.74 |
| EducationalAttainment | DNase | Fetal Brain Female | 3.05 |
| EducationalAttainment | DNase-gapped | Cortex derived primary cultured neurospheres | 4.27 |
| EducationalAttainment | DNase-narrow | Fetal Brain Female | 3.07 |
| EducationalAttainment | FUN-LDA | Fetal Brain Female | 5.84 |
| EducationalAttainment | GenoSkyline | Brain Dorsolateral Prefrontal Cortex | 3.61 |
| EducationalAttainment | IDEAS | Fetal Brain Female | 7.32 |
| EducationalAttainment | Segway | Fetal Brain Male | 5.55 |
| Epilepsy | ChromHMM | Brain Angular Gyrus | 2.91 |
| Epilepsy | DNase | Dnd41 TCell Leukemia Cell Line | 0.99 |
| Epilepsy | DNase-gapped | Brain Hippocampus Middle | 2.36 |
| Epilepsy | DNase-narrow | Fetal Thymus | 1.85 |
| Epilepsy | FUN-LDA | Brain Anterior Caudate | 4.11 |
| Epilepsy | GenoSkyline | Brain Inferior Temporal Lobe | 3.35 |
| Epilepsy | IDEAS | Brain Angular Gyrus | 4.40 |
| Epilepsy | Segway | Brain Angular Gyrus | 4.51 |
| EverSmoked | ChromHMM | Primary T cells effector/memory enriched from peripheral blood | 2.15 |
| EverSmoked | DNase | Brain Inferior Temporal Lobe | 0.61 |
| EverSmoked | DNase-gapped | Brain Inferior Temporal Lobe | 1.31 |
| EverSmoked | DNase-narrow | Primary hematopoietic stem cells | 0.78 |
| EverSmoked | FUN-LDA | Brain Inferior Temporal Lobe | 2.68 |
| EverSmoked | GenoSkyline | Brain Inferior Temporal Lobe | 2.94 |
| EverSmoked | IDEAS | Brain Angular Gyrus | 3.66 |
| EverSmoked | Segway | Brain Inferior Temporal Lobe | 4.16 |
| FastingGlucose | ChromHMM | Pancreatic Islets | 1.44 |
| FastingGlucose | DNase | Fetal Intestine Small | 1.03 |
| FastingGlucose | DNase-gapped | Pancreatic Islets | 2.03 |
| FastingGlucose | DNase-narrow | iPS-15b Cells | 1.60 |
| FastingGlucose | FUN-LDA | Pancreatic Islets | 1.45 |
| FastingGlucose | GenoSkyline | H9 Cells | 2.29 |
| FastingGlucose | IDEAS | Pancreatic Islets | 3.65 |
| FastingGlucose | Segway | Pancreatic Islets | 3.85 |
| HDL | ChromHMM | Primary monocytes from peripheral blood | 2.72 |
| HDL | DNase | Liver | 3.94 |
| HDL | DNase-gapped | Adipose Nuclei | 5.15 |
| HDL | DNase-narrow | Adipose Nuclei | 4.37 |
| HDL | FUN-LDA | Liver | 4.73 |
| HDL | GenoSkyline | Liver | 3.67 |
| HDL | IDEAS | Adipose Nuclei | 5.63 |
| HDL | Segway | Liver | 4.28 |
| Height | ChromHMM | Mesenchymal Stem Cell Derived Chondrocyte Cultured Cells | 5.55 |
| Height | DNase | Mesenchymal Stem Cell Derived Chondrocyte Cultured Cells | 4.45 |
| Height | DNase-gapped | Mesenchymal Stem Cell Derived Chondrocyte Cultured Cells | 9.99 |
| Height | DNase-narrow | Mesenchymal Stem Cell Derived Chondrocyte Cultured Cells | 10.81 |
| Height | FUN-LDA | Mesenchymal Stem Cell Derived Chondrocyte Cultured Cells | 12.28 |
| Height | GenoSkyline | Mesenchymal Stem Cell Derived Chondrocyte Cultured Cells | 11.31 |
| Height | IDEAS | Mesenchymal Stem Cell Derived Chondrocyte Cultured Cells | 14.59 |
| Height | Segway | Mesenchymal Stem Cell Derived Chondrocyte Cultured Cells | 13.40 |

**Table s7:**
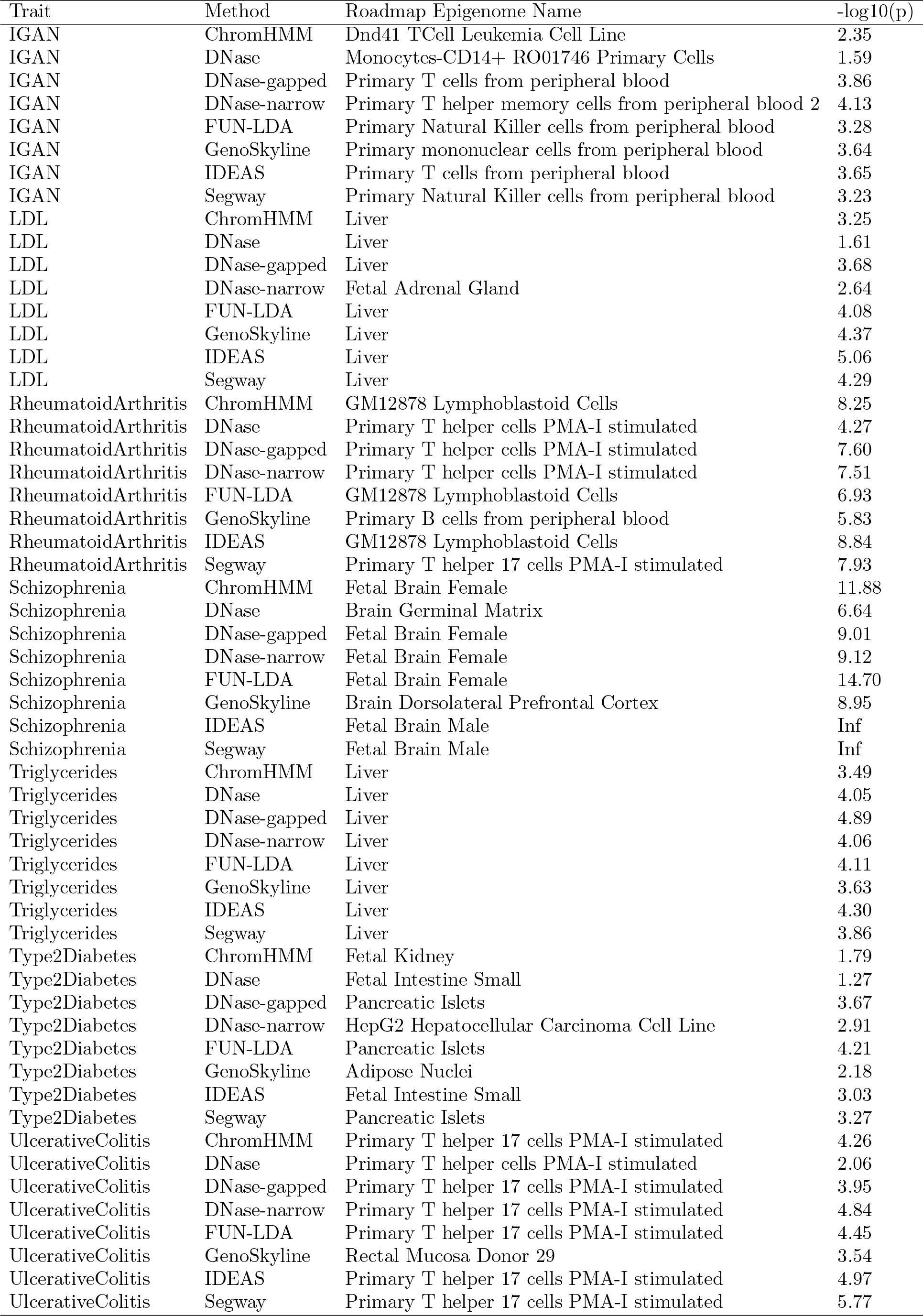
Results from stratified LD score regression for the different methods (part 3).

| Trait | Method | Roadmap Epigenome Name | -log10(p) |
| --- | --- | --- | --- |
| IGAN | ChromHMM | Dnd41 TCell Leukemia Cell Line | 2.35 |
| IGAN | DNase | Monocytes-CD14+ RO01746 Primary Cells | 1.59 |
| IGAN | DNase-gapped | Primary T cells from peripheral blood | 3.86 |
| IGAN | DNase-narrow | Primary T helper memory cells from peripheral blood 2 | 4.13 |
| IGAN | FUN-LDA | Primary Natural Killer cells from peripheral blood | 3.28 |
| IGAN | GenoSkyline | Primary mononuclear cells from peripheral blood | 3.64 |
| IGAN | IDEAS | Primary T cells from peripheral blood | 3.65 |
| IGAN | Segway | Primary Natural Killer cells from peripheral blood | 3.23 |
| LDL | ChromHMM | Liver | 3.25 |
| LDL | DNase | Liver | 1.61 |
| LDL | DNase-gapped | Liver | 3.68 |
| LDL | DNase-narrow | Fetal Adrenal Gland | 2.64 |
| LDL | FUN-LDA | Liver | 4.08 |
| LDL | GenoSkyline | Liver | 4.37 |
| LDL | IDEAS | Liver | 5.06 |
| LDL | Segway | Liver | 4.29 |
| RheumatoidArthritis | ChromHMM | GM12878 Lymphoblastoid Cells | 8.25 |
| RheumatoidArthritis | DNase | Primary T helper cells PMA-I stimulated | 4.27 |
| RheumatoidArthritis | DNase-gapped | Primary T helper cells PMA-I stimulated | 7.60 |
| RheumatoidArthritis | DNase-narrow | Primary T helper cells PMA-I stimulated | 7.51 |
| RheumatoidArthritis | FUN-LDA | GM12878 Lymphoblastoid Cells | 6.93 |
| RheumatoidArthritis | GenoSkyline | Primary B cells from peripheral blood | 5.83 |
| RheumatoidArthritis | IDEAS | GM12878 Lymphoblastoid Cells | 8.84 |
| RheumatoidArthritis | Segway | Primary T helper 17 cells PMA-I stimulated | 7.93 |
| Schizophrenia | ChromHMM | Fetal Brain Female | 11.88 |
| Schizophrenia | DNase | Brain Germinal Matrix | 6.64 |
| Schizophrenia | DNase-gapped | Fetal Brain Female | 9.01 |
| Schizophrenia | DNase-narrow | Fetal Brain Female | 9.12 |
| Schizophrenia | FUN-LDA | Fetal Brain Female | 14.70 |
| Schizophrenia | GenoSkyline | Brain Dorsolateral Prefrontal Cortex | 8.95 |
| Schizophrenia | IDEAS | Fetal Brain Male | Inf |
| Schizophrenia | Segway | Fetal Brain Male | Inf |
| Triglycerides | ChromHMM | Liver | 3.49 |
| Triglycerides | DNase | Liver | 4.05 |
| Triglycerides | DNase-gapped | Liver | 4.89 |
| Triglycerides | DNase-narrow | Liver | 4.06 |
| Triglycerides | FUN-LDA | Liver | 4.11 |
| Triglycerides | GenoSkyline | Liver | 3.63 |
| Triglycerides | IDEAS | Liver | 4.30 |
| Triglycerides | Segway | Liver | 3.86 |
| Type2Diabetes | ChromHMM | Fetal Kidney | 1.79 |
| Type2Diabetes | DNase | Fetal Intestine Small | 1.27 |
| Type2Diabetes | DNase-gapped | Pancreatic Islets | 3.67 |
| Type2Diabetes | DNase-narrow | HepG2 Hepatocellular Carcinoma Cell Line | 2.91 |
| Type2Diabetes | FUN-LDA | Pancreatic Islets | 4.21 |
| Type2Diabetes | GenoSkyline | Adipose Nuclei | 2.18 |
| Type2Diabetes | IDEAS | Fetal Intestine Small | 3.03 |
| Type2Diabetes | Segway | Pancreatic Islets | 3.27 |
| UlcerativeColitis | ChromHMM | Primary T helper 17 cells PMA-I stimulated | 4.26 |
| UlcerativeColitis | DNase | Primary T helper cells PMA-I stimulated | 2.06 |
| UlcerativeColitis | DNase-gapped | Primary T helper 17 cells PMA-I stimulated | 3.95 |
| UlcerativeColitis | DNase-narrow | Primary T helper 17 cells PMA-I stimulated | 4.84 |
| UlcerativeColitis | FUN-LDA | Primary T helper 17 cells PMA-I stimulated | 4.45 |
| UlcerativeColitis | GenoSkyline | Rectal Mucosa Donor 29 | 3.54 |
| UlcerativeColitis | IDEAS | Primary T helper 17 cells PMA-I stimulated | 4.97 |
| UlcerativeColitis | Segway | Primary T helper 17 cells PMA-I stimulated | 5.77 |

**Table s8:** For eight SNPs selected from literature, the tissue or cell type in the original study and the closest tissue in Roadmap that we selected are given.

| SNP | Tissue in Functional Study | Selected Roadmap Tissue |
| --- | --- | --- |
| rs6801957 | murine heart tissue | E104 - Right Atrium |
| rs12821256 | cultured human keratinocytes | E127 - NHEK-Epidermal Keratinocyte Primary Cells |
| rs12350739 | skin epidermal samples/melanocyte cell lines | E059 - Foreskin Melanocyte Primary Cells skin01 |
| rs12740374 | primary hepatocytes | E066 - Liver |
| rs356168 | hIPSC-derived neurons | E007 - H1 Derived Neuronal Progenitor Cultured Cells |
| rs2473307 | human neuronal cell line | E007 - H1 Derived Neuronal Progenitor Cultured Cells |
| rs227727 | human embryonic oral epithelial cells | E119 - HMEC Mammary Epithelial Primary Cells |
| rs144361550 | primary melanocytes | E059 - Foreskin Melanocyte Primary Cells skin01 |

**Table s9:** AUC for various integrative methods vs. individual epigenetic annotations using MPRA validated variants.

| Method | Type | emVars<br>E116 | Regulatory motifs<br>E118 | Regulatory motifs<br>E123 |
| --- | --- | --- | --- | --- |
| FUN-LDA | Integrative | 0.709 | 0.694 | 0.646 |
| GenoSkyline |  | 0.674 | 0.630 | 0.619 |
| ChromHMM |  | 0.668 | 0.608 | 0.634 |
| Segway |  | 0.624 | 0.618 | 0.585 |
| IDEAS |  | 0.621 | 0.546 | 0.615 |
| DNase | Single annotation | 0.722 | 0.719 | 0.654 |
| DNase-narrow |  | 0.629 | 0.561 | 0.524 |
| DNase-gapped |  | 0.653 | 0.550 | 0.565 |
| H3K27ac |  | 0.677 | 0.556 | 0.597 |
| H3K4me1 |  | 0.664 | 0.545 | 0.578 |
| H3K4me3 |  | 0.692 | 0.535 | 0.602 |
| H3K9ac |  | 0.670 | 0.549 | 0.615 |

**Table s10:** AUC for the segmentation methods ChromHMM, Segway and IDEAS state combinations with maximum AUC using the MPRA validated variants. Note that the selection of the best state combination is based on combining the variants from all three MPRA datasets in Section 2.4.

| Method | Type | State | States in 'functional' group | emVars<br>E116 | Reg. motifs<br>E118 | Reg. motifs<br>E123 |
| --- | --- | --- | --- | --- | --- | --- |
| FUN-LDA | Selected |  | 1_ActiveEnhancers, 2_ActivePromoters | 0.709 | 0.694 | 0.646 |
| ChromHMM | Best |  | 1_TssA, 2_PromU, 9_TxReg, 13_EnhA1<br>14_EnhA2, 16_EnhW1, 22_PromP | 0.670 | 0.619 | 0.661 |
|  | Selected |  | 1_TssA, 2_PromU, 3_PromD1, 4_PromD2<br>13_EnhA1, 14_EnhA2, 15_EnhAF | 0.668 | 0.608 | 0.634 |
| Segway | Best |  | Bivalent, RegPermissive, Enhancer, Promoter | 0.650 | 0.591 | 0.630 |
|  | Selected |  | Enhancer, Promoter | 0.624 | 0.618 | 0.585 |
| IDEAS | Best |  | 4_Enh, 8_TssAFlnk, 6_EnhG, 10_TssA<br>19_Enh/ReprPC, 11_EnhBiv, 15_TssBiv, 14_TssWk, 17_EnhGA | 0.635 | 0.544 | 0.614 |
|  | Selected |  | 4_Enh, 6_EnhG, 8_TssAFlnk, 10_TssA, 14_TssWk, 17_EnhGA | 0.621 | 0.546 | 0.615 |

**Table s11:** Grouping of Roadmap tissues into 10 tissue types.

| Epigenome.ID | Type | Epigenome.ID | Type |
| --- | --- | --- | --- |
| E022 | Stem cell | E117 | Connective tissue |
| E007 | Stem cell | E028 | Connective tissue |
| E004 | Stem cell | E057 | Connective tissue |
| E002 | Stem cell | E058 | Connective tissue |
| E021 | Stem cell | E119 | Connective tissue |
| E009 | Stem cell | E127 | Connective tissue |
| E010 | Stem cell | E071 | Brain |
| E001 | Stem cell | E074 | Brain |
| E015 | Stem cell | E073 | Brain |
| E018 | Stem cell | E068 | Brain |
| E016 | Stem cell | E067 | Brain |
| E020 | Stem cell | E069 | Brain |
| E014 | Stem cell | E072 | Brain |
| E019 | Stem cell | E027 | Internal organs |
| E024 | Stem cell | E059 | Internal organs |
| E008 | Stem cell | E061 | Internal organs |
| E003 | Stem cell | E065 | Internal organs |
| E012 | Stem cell | E097 | Internal organs |
| E011 | Stem cell | E086 | Internal organs |
| E115 | Blood | E087 | Internal organs |
| E123 | Blood | E100 | Internal organs |
| E030 | Blood | E105 | Internal organs |
| E029 | Blood | E104 | Internal organs |
| E124 | Blood | E095 | Internal organs |
| E035 | Blood | E096 | Internal organs |
| E036 | Blood | E113 | Internal organs |
| E051 | Blood | E079 | Internal organs |
| E050 | Blood | E094 | Internal organs |
| E034 | Blood | E098 | Internal organs |
| E046 | Blood | E081 | Fetal brain |
| E041 | Blood | E070 | Fetal brain |
| E047 | Blood | E082 | Fetal brain |
| E048 | Blood | E054 | Fetal brain |
| E038 | Blood | E053 | Fetal brain |
| E045 | Blood | E005 | Fetal tissue 1 |
| E044 | Blood | E099 | Fetal tissue 1 |
| E043 | Blood | E013 | Fetal tissue 1 |
| E039 | Blood | E006 | Fetal tissue 1 |
| E042 | Blood | E083 | Fetal tissue 1 |
| E040 | Blood | E108 | Muscle |
| E037 | Blood | E107 | Muscle |
| E112 | Blood | E063 | Muscle |
| E093 | Blood | E078 | Muscle |
| E062 | Blood | E103 | Muscle |
| E033 | Blood | E076 | Muscle |
| E116 | Blood | E111 | Muscle |
| E031 | Blood | E091 | Fetal tissue 2 |
| E032 | Blood | E092 | Fetal tissue 2 |
| E122 | Connective tissue | E089 | Fetal tissue 2 |
| E120 | Connective tissue | E090 | Fetal tissue 2 |
| E121 | Connective tissue | E088 | Fetal tissue 2 |
| E025 | Connective tissue | E080 | Fetal tissue 2 |
| E023 | Connective tissue | E066 | GI |
| E049 | Connective tissue | E110 | GI |
| E026 | Connective tissue | E109 | GI |
| E129 | Connective tissue | E106 | GI |
| E126 | Connective tissue | E075 | GI |
| E052 | Connective tissue | E077 | GI |
| E125 | Connective tissue | E101 | GI |
| E055 | Connective tissue | E102 | GI |
| E056 | Connective tissue | E118 | GI |
| E017 | Connective tissue | E085 | GI |
| E128 | Connective tissue | E084 | GI |
| E114 | Connective tissue |  |  |

## References

Lindblad-Toh K et al. (2011) A high-resolution map of human evolutionary constraint using 29 mammals. Nature 478: 476–482.

Khurana E et al. (2013) Integrative annotation of variants from 1092 humans: application to cancer genomics. Science 342: 1235587.

ENCODE Project Consortium (2012) An integrated encyclopedia of DNA elements in the human genome. Nature 489: 57–74.

Altshuler D, Daly MJ, Lander ES (2008) Genetic mapping in human disease. Science 322: 881–888.

Khurana E, Fu Y, Chakravarty D, Demichelis F, Rubin MA, Gerstein M (2016) Role of non-coding sequence variants in cancer. Nat Rev Genet 17: 93–108.

Kellis M et al. (2014) Defining functional DNA elements in the human genome. Proc Natl Acad Sci USA 111: 6131–6138.

Petrovski S, Wang Q, Heinzen EL, Allen AS, Goldstein DB (2013) Genic intolerance to functional variation and the interpretation of personal genomes. PLoS Genet 9: e1003709.

Roadmap Epigenomics Consortium (2015) Integrative analysis of 111 reference human epigenomes. Nature 518: 317–330.

Kircher M, Witten DM, Jain P, O’Roak BJ, Cooper GM, Shendure J (2014) A general framework for estimating the relative pathogenicity of human genetic variants. Nat Genet 46: 310–315.

Fu Y, Liu Z, Lu S, Bedford J, Mu X, Yip K, Khurana E, Gerstein M (2014) FunSeq2: A framework for prioritizing noncoding regulatory variants in cancer. Genome Biology 15: 480

Ionita-Laza I, McCallum K, Xu B, Buxbaum JD (2016) A spectral approach integrating functional genomic annotations for coding and noncoding variants. Nat Genet 48: 214–220.

Bannister AJ, Kouzarides T (2011) Regulation of chromatin by histone modifications. Cell Res. 21(3): 381–395.

Friedman N, Rando OJ (2015) Epigenomics and the structure of the living genome Genome Res 25: 1482–1490.

Ernst J, Kellis M (2012) ChromHMM: automating chromatin-state discovery and characterization. Nature Methods 9: 215–216.

Ernst J, Kellis M (2015) Large-scale imputation of epigenomic datasets for systematic annotation of diverse human tissues. Nat Biotechnol 33: 364–376.

Hoffman MM, Buske OJ, Wang J, Weng Z, Bilmes J, Noble WS (2012) Unsupervised pattern discovery in human chromatin structure through genomic segmentation. Nat Methods 9: 473–476.

Zacher B, Michel M, Schwalb B, Cramer P, Tresch A, Gagneur J (2017) Accurate Promoter and Enhancer Identification in 127 ENCODE and Roadmap Epigenomics Cell Types and Tissues by GenoSTAN. PLoS One 12: e0169249.

Mammana A, Chung HR (2015) Chromatin segmentation based on a probabilistic model for read counts explains a large portion of the epigenome. Genome Biol 16: 151.

Biesinger J, Wang Y, Xie X (2013) Discovering and mapping chromatin states using a tree hidden Markov model. BMC Bioinformatics Suppl 5: S4.

Zhang Y, An L, Yue F, Hardison RC (2016) Jointly characterizing epigenetic dynamics across multiple human cell types. Nucleic Acids Res 44: 6721–6731.

Zhang Y, Hardison RC (2017) Accurate and Reproducible Functional Maps in 127 Human Cell Types via 2D Genome Segmentation. BioRxiv preprint doi: http://dx.doi.org/10.1101/118752.

Song J, Chen KC (2015) Spectacle: fast chromatin state annotation using spectral learning. Genome Biol 16: 33.

Lu Q, Powles RL, Wang Q, He BJ, Zhao H (2016) Integrative Tissue-Specific Functional Annotations in the Human Genome Provide Novel Insights on Many Complex Traits and Improve Signal Prioritization in Genome Wide Association Studies. PLoS Genet 12: e1005947.

Blei DM, Ng AY, Jordan MI (2003) Latent Dirichlet Allocation. Journal of Machine Learning Research 3: 9931022.

Ramsey S et al. (2010) Genome-wide histone acetylation data improve prediction of mammalian transcription factor binding sites. Bioinformatics 26: 2071–2075.

Heintzman ND et al. (2009) Histone modifications at human enhancers reflect global cell-type-specific gene expression. Nature 459: 108–112.

The GTEx Consortium (2015) Science. 348: 648–660.

Brown AA, Vi?nuela A, Delaneau O, Spector T, Small K, Dermitzakis ET (2016) Predicting causal variants affecting expression using whole-genome sequence and RNA-seq from multiple human tissues. http://www.biorxiv.org/content/biorxiv/early/2016/11/21/088872.full.pdf

Finucane HK et al. (2015) Partitioning heritability by functional annotation using genome-wide association summary statistics. Nat Genet 47: 1228–1235.

Perry JR et al. (2014) Parent-of-origin-specific allelic associations among 106 genomic loci for age at menarche. Nature 514: 92–97.

Betz RC et al. (2015) Genome-wide meta-analysis in alopecia areata resolves HLA associations and reveals two new susceptibility loci. Nat Commun 6: 5966.

Lambert JC et al. (2013) Meta-analysis of 74,046 individuals identifies 11 new susceptibility loci for Alzheimer’s disease. Nat Genet 45: 1452–1458.

Cross-Disorder Group of the Psychiatric Genomics Consortium (2013) Identification of risk loci with shared effects on five major psychiatric disorders: a genome-wide analysis. Lancet 381: 1371–1379.

Psychiatric GWAS Consortium Bipolar Disorder Working Group (2011) Large-scale genome-wide association analysis of bipolar disorder identifies a new susceptibility locus near ODZ4. Nat Genet 43: 977–983.

Speliotes EK et al. (2010) Association analyses of 249,796 individuals reveal 18 new loci associated with body mass index. Nat Genet 42: 937–948.

Schunkert H et al. (2011) Large-scale association analysis identifies 13 new susceptibility loci for coronary artery disease. Nat Genet 43: 333–338.

Jostins L et al. (2012) Host-microbe interactions have shaped the genetic architecture of inflammatory bowel disease. Nature 491: 119–124.

Petukhova L, Christiano AM (2016) Functional Interpretation of Genome-Wide Association Study Evidence in Alopecia Areata. The Journal of investigative dermatology 136: 314–317.

Xing L et al. (2014) Alopecia areata is driven by cytotoxic T lymphocytes and is reversed by JAK inhibition. Nature medicine 20: 1043–1049.

Yokoyama JS et al. (2016) Association Between Genetic Traits for Immune-Mediated Diseases and Alzheimer Disease. JAMA Neurol 73: 691–697.

Rietveld CA et al. (2013) GWAS of 126,559 individuals identifies genetic variants associated with educational attainment. Science 314: 1467–1471.

International League Against Epilepsy Consortium on Complex Epilepsies (2014) Genetic determinants of common epilepsies: a meta-analysis of genome-wide association studies. Lancet Neurol 13: 893–903.

Tobacco and Genetics Consortium (2010) Genome-wide meta-analyses identify multiple loci associated with smoking behavior. Nat Genet 42:441–447.

Manning AK et al. (2012) A genome-wide approach accounting for body mass index identifies genetic variants influencing fasting glycemic traits and insulin resistance. Nat Genet 44: 659–669.

Teslovich TM et al. (2010) Biological, clinical and population relevance of 95 loci for blood lipids. Nature 466: 707–713.

Kiryluk K et al. (2014) Discovery of new risk loci for IgA nephropathy implicates genes involved in immunity against intestinal pathogens. Nat Genet 46: 1187–1196.

Okada Y et al. (2014) Genetics of rheumatoid arthritis contributes to biology and drug discovery. Nature 506: 376–381.

Schizophrenia Working Group of the Psychiatric Genomics Consortium (2014) Biological insights from 108 schizophrenia-associated genetic loci. Nature 511: 421–427.

Morris AP et al. (2012) Large-scale association analysis provides insights into the genetic architecture and pathophysiology of type 2 diabetes. Nat Genet 44: 981–990.

Lango AH et al. (2010) Hundreds of variants clustered in genomic loci and biological pathways affect human height. Nature 467: 832–838.

Locke AE et al. (2015) Genetic studies of body mass index yield new insights for obesity biology. Nature 518: 197–206.

Magga J et al. (2012) Production of monocytic cells from bone marrow stem cells: therapeutic usage in Alzheimer’s disease. J Cell Mol Med 16: 1060–1073.

Gjoneska E, Pfenning AR, Mathys H, Quon G, Kundaje A, Tsai LH, Kellis M (2015) Conserved epigenomic signals in mice and humans reveal immune basis of Alzheimer’s disease. Nature 518: 365–369.

Bulik-Sullivan B et al. An atlas of genetic correlations across human diseases and traits. Nat Genet 47: 1236–1241.

Jefferson AL et al. (2015) Low cardiac index is associated with incident dementia and Alzheimer disease: the Framingham Heart Study. Circulation 131: 1333–1339.

van den Boogaard M et al. (2014) A common genetic variant within scn10a modulates cardiac scn5a expression. J Clin Invest 124: 1844–1852.

Guenther CA, Tasic B, Luo L, Bedell MA, Kingsley DM (2014) A molecular basis for classic blond hair color in europeans. Nat Genet 46: 748–752.

Visser M, Palstra RJ, Kayser M (2014) Human skin color is influenced by an intergenic dna polymorphism regulating transcription of the nearby bnc2 pigmentation gene. Hum Mol Genet 23: 5750–5562.

Musunuru K et al. (2010) From noncoding variant to phenotype via SORT1 at the 1p13 cholesterol locus. Nature 466: 714–719.

Soldner F et al. (2016) Parkinson-associated risk variant in distal enhancer of ?-synuclein modulates target gene expression. Nature 533: 95–99.

Gilks WP, Hill M, Gill M, Donohoe G, Corvin AP, Morris DW (2012) Functional investigation of a schizophrenia gwas signal at the cdc42 gene. World J Biol Psychiatry 13: 550–554.

Leslie EJ et al. (2015) Identification of functional variants for cleft lip with or without cleft palate in or near PAX7, FGFR2, and NOG by targeted sequencing of GWAS loci. Am J Hum Genet 96: 397–411.

Choi J et al. (2017) A common intronic variant of PARP1 confers melanoma risk and mediates melanocyte growth via regulation of MITF. Nat Genet Epub ahead of print

Tewhey R et al. (2016) Direct identification of hundreds of expression-modulating variants using a multiplexed reporter assay. Cell 165: 1519–1529.

Kheradpour P, Ernst J, Melnikov A, Rogov P, Wang L, Zhang X, Alston J, Mikkelsen TS, Kellis M (2013) Systematic dissection of regulatory motifs in 2000 predicted human enhancers using a massively parallel reporter assay. Genome Res 23: 800–811.

Pennacchio LA et al. (2006) In vivo enhancer analysis of human conserved non-coding sequences. Nature 444: 499–502.

Lee S, Wu MC, Lin X (2012) Optimal tests for rare variant effects in sequencing association studies. Biostatistics 13: 762–775.

Ionita-Laza I, Capanu M, De Rubeis S, McCallum K, Buxbaum JD (2014) Identification of rare causal variants in sequence-based studies: methods and applications to VPS13B, a gene involved in Cohen syndrome and autism. PLoS Genet 10: e1004729.

Kichaev G, Yang WY, Lindstrom S, Hormozdiari F, Eskin E, Price AL, Kraft P, Pasaniuc B (2014) Integrating functional data to prioritize causal variants in statistical fine-mapping studies. PLoS Genet 10: e1004722.

Silverman BW (1986) Density Estimation for Statistics and Data Analysis, Chapman & Hall, London

Hagai Attias (1999) Inferring parameters and structure of latent variable models by variational bayes. Proceedings of the Fifteenth conference on Uncertainty in artificial intelligence, Morgan Kaufmann Publishers Inc., pp. 21–30.

Libbrecht MW, Rodriguez O, Weng Z, Hoffman M, Bilmes JA, Noble WS (2017) A unified encyclopedia of human functional DNA elements through fully automated annotation of 164 human cell types. doi: https://doi.org/10.1101/086025

Xiaoyi G, Starmer J, Martin ER (2008) A multiple testing correction method for genetic association studies using correlated single nucleotide polymorphisms. Genetic Epidemiology 32: 361–369.

## References

1,000 Genomes Project Consortium (2012) An integrated map of genetic variation from 1,092 human genomes. Nature 491: 56–65.

Blei DM, Ng AY, Jordan MI (2003) Latent Dirichlet Allocation. Journal of Machine Learning Research 3: 993–1022.

